# Consensus sequence design as a general strategy to create hyperstable, biologically active proteins

**DOI:** 10.1101/466391

**Authors:** Matt Sternke, Katherine W. Tripp, Doug Barrick

## Abstract

Consensus sequence design offers a promising strategy for designing proteins of high stability while retaining biological activity since it draws upon an evolutionary history in which residues important for both stability and function are likely to be conserved. Although there have been several reports of successful consensus design of individual targets, it is unclear from these anecdotal studies how often this approach succeeds, and how often it fails. Here, we attempt to assess generality by designing consensus sequences for a set of six protein families with a range of chain-lengths, structures, and activities. We characterize the resulting consensus proteins for stability, structure, and biological activities in an unbiased way. We find that all six consensus proteins adopt cooperatively folded structures in solution. Strikingly, four out of six of these consensus proteins show increased thermodynamic stability over naturally-occurring homologues. Each consensus protein tested for function maintained at least partial biological activity. Though peptide binding affinity by a consensus-designed SH3 is rather low, K_m_ values for consensus enzymes are similar to values from extant homologues. Though consensus enzymes are slower than extant homologues at low temperature, they are faster than some thermophilic enzymes at high temperature. An analysis of sequence properties shows consensus proteins to be enriched in charged residues, and rarified in uncharged polar residues. Sequence differences between consensus and extant homologues are predominantly located at weakly conserved surface residues, highlighting the importance of these residues in the success of the consensus strategy.

**Significance Statement:** A major goal of protein design is to create proteins that have high stability and biological activity. Drawing on evolutionary information encoded within extant protein sequences, consensus sequence design has produced several successes in achieving this goal. Here we explore the generality with which consensus design can be used to enhance protein stability and maintain biological activity. By designing and characterizing consensus sequences for six unrelated protein families, we find that consensus design shows high success rates in creating well-folded, hyperstable proteins that retain biological activities. Remarkably, many of these consensus proteins show higher stabilities than naturally-occurring sequences of their respective protein families. Our study highlights the utility of consensus sequence design and informs the mechanisms by which it works.

## Introduction

Exploiting the fundamental roles of proteins in biological signaling, catalysis, and mechanics, protein design offers a promising route to create and optimize biomolecules for medical, industrial, and biotechnological purposes (1–3). Many different strategies have been applied to designing proteins, including physics-based (4), structure-based (5, 6), and directed evolution-based approaches (7). While these strategies have generated proteins with high stability, implementation of the design strategies is often complex and success rates can be low (8–11). Though directed evolution can be functionally directed, *de novo* design strategies typically focus primarily on structure. Introducing specific activity into *de novo* designed proteins is a significant challenge (8, 10).

Another strategy that has shown success in increasing thermodynamic stability of natural protein folds is consensus sequence design (12). For this design strategy, “consensus” residues are identified as the residues with the highest frequency at individual positions in a multiple sequence alignment (MSA) of extant sequences from a given protein family. The consensus strategy draws upon the hundreds of millions of years of sequence evolution by random mutation and natural selection that is encoded within the extant sequence distribution, with the idea that the relative frequencies of residues at a given position reflect the “relative importance” of each residue at that position for some biological attribute. As long as the importance at each position is largely independent of residues at other positions, a consensus residue at a given position should optimize stability, activity, and/or other properties that allow the protein to function in its biological context and ultimately contribute to organismal fitness^†^. By averaging over many sequences that share similar structure and function, the consensus design approach has the potential to produce proteins with high levels of thermodynamic stability and biological activity, since both attributes are likely to lead to residue conservation.

There are two experimental approaches that have been used to examine the effectiveness of consensus information in protein design: point-substitution and “wholesale” substitution. In the first approach, single residues in a well-behaved protein that differ from the consensus are substituted with the consensus residue (13–17). In these studies, about half of the consensus point substitutions examined are stabilizing, but the other half are destabilizing. Although this frequency of stabilizing mutations is significantly higher than an estimated frequency around one in 10^3^ for random mutations (13, 18, 19), it suggests that combining individual consensus mutations may give minimal net increase in stability since stabilizing substitutions would be offset by destabilizing substitutions.

The “wholesale” approach does just this, combining all substitutions toward consensus into a single consensus polypeptide composed of the most frequent amino acid at each position in sequence. By making a large number of substitutions at once, wholesale consensus substitution may collectively combine the incremental effects from the individual substitutions as well as non-additive effects arising from the substitution of each residue into the novel background of the consensus protein (20, 21). The stabilities of several globular proteins and several repeat proteins have been increased using this approach (22–31). An increase in thermodynamic stability is seen in most (but not all) cases, but effects on biological activity are variable. In a recent study, our lab characterized a consensus-designed homeodomain sequence that showed a large increase in both thermodynamic stability and DNA-binding affinity (32). Unlike the point-substitution approach, where both stabilizing and destabilizing substitutions are reported, the success rate of the wholesale approach is not easy to determine from the literature, where publications present single cases of success, whereas failures are not likely to be published. One study of TIM barrels reported a few poorly behaved consensus designs that were then optimized to generate a folded, active protein (24). Although this study highlighted some limitations, it would not likely have been published if it did not end with success.

Here, we address these issues by applying the consensus sequence design strategy to a set of six taxonomically diverse protein families with different folds and functions (Figure 1). We chose the three single domain families including the N-terminal domain of ribosomal protein L9 (NTL9), the SH3 domain, and the SH2 domain, and the three multi-domain protein families including dihydrofolate reductase (DHFR), adenylate kinase (AK), and phosphoglycerate kinase (PGK). We characterized these six consensus proteins in terms of structure, stability, and function. We find that consensus sequences for all six protein families are quite soluble, and adopt the native folds of their respective families. Strikingly, four of the six consensus proteins show increased thermodynamic stability compared to naturally-occurring homologues; the other two consensus proteins show stabilities comparable to natural homologues. All consensus proteins assayed for biological activity retain their expected activities, including molecular recognition and enzymatic catalysis. An advantage of this multi-target comparison is that it allows us to examine sequence features of consensus-designed proteins and relate them to one another and to naturally occurring homologues. This sequence analysis shows that consensus proteins are enriched in charged residues, are depleted in polar uncharged residues, and highlights the importance of weakly conserved surface residues in enhancing stability through the consensus design strategy.

**Figure 1.**
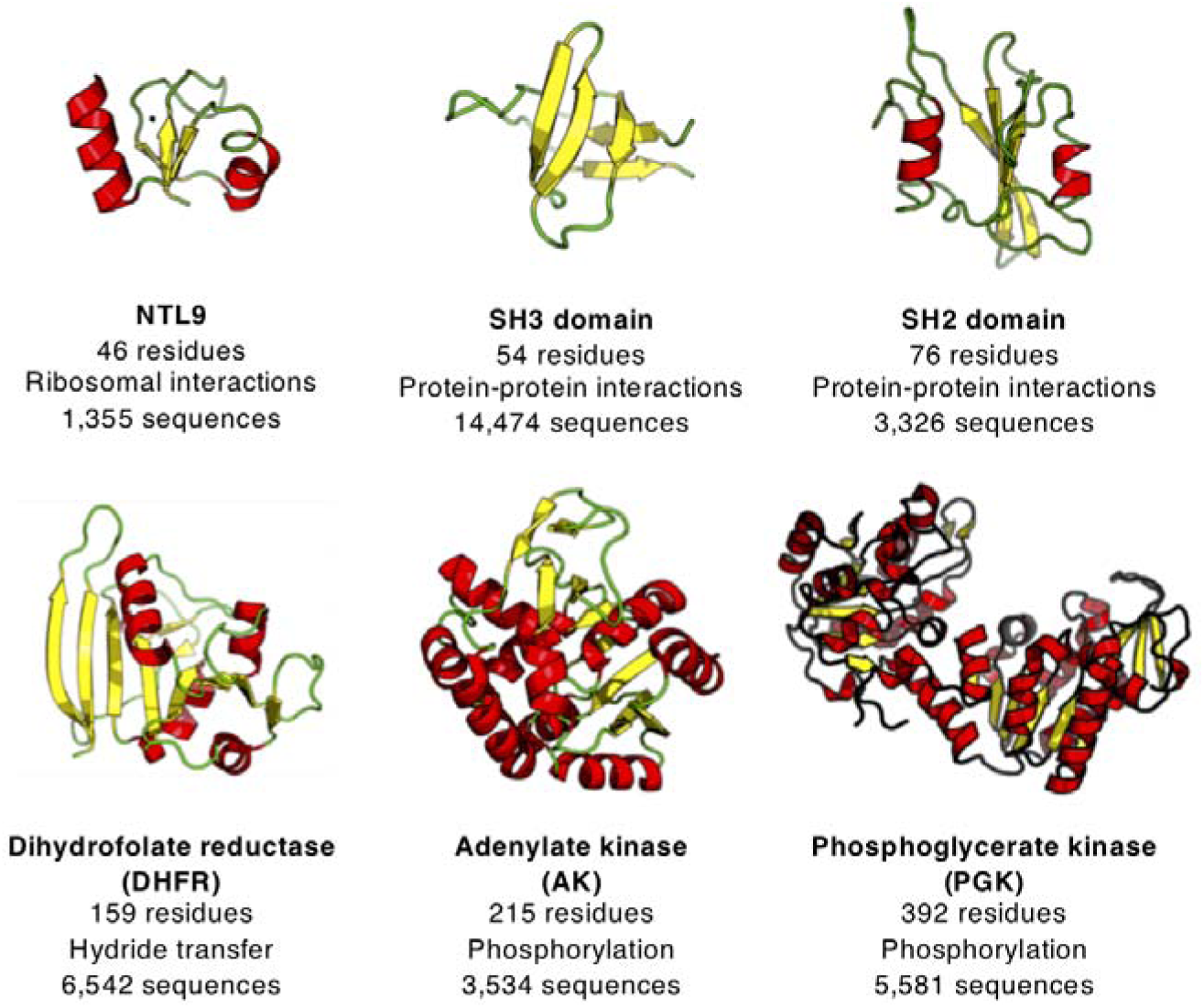
Targets for consensus design. A representative structure of an extant sequence is shown for each target family (NTL9: 2HBB, SH3: 1LKK, SH2: 4U1P, DHFR: 5DFR, AK: 1ANK, PGK: 1PHP). Length of consensus sequence, biological function, and number of sequences used in final multiple sequence alignment are noted.

## Results

### Consensus sequence generation

The six families targeted for consensus design are shown in Figure 1. To generate and refine multiple sequence alignments from these families, we surveyed the Pfam (33), SMART (34), and InterPro (35) databases to obtain sets of aligned sequences for each family. For each family, we used the sequence set that maximized both the number of sequences and quality of the multiple sequence alignment (SI Appendix, Table S1). For five of the six protein families, we did not limit sequences using any structural or phylogenetic information. For AK, we selected sequences that contained a "long type" LID domain containing a 27-residue insertion found primarily in bacterial sequences (36). The number of sequences in these initial sequence sets ranged from around 2,000 up to 55,000 (SI Appendix, Table S1).

To improve alignments and limit bias from copies of identical (or nearly identical) sequences, we removed sequences that deviated significantly (±30%) in length from the median length value, and removed sequences that shared greater than 90% sequence identity with another sequence. This curation reduced the size of sequence sets by an average of 65% (ranging from 30% to 80%; SI Appendix, Table S1). Final sequence sets ranged from 1,355 (NTL9) to 14,474 (SH3) sequences, and showed broad ranges of sequence variability (quantified by average pairwise identities among sequences in the set) and phylogenetic distribution (SI Appendix, Table S1).

MSAs showed varying levels of conservation across positions in sequence (SI Appendix, Figure S1). Curated sequence sets were aligned, and consensus sequences were determined from the most probable residue at each position (SI Appendix, Table S2). Consensus sequences differ substantially from aligned sequences from which they are derived. The maximum identity between consensus sequences and the most similar extant sequence in the corresponding alignment ranges from 63% for SH2 to 80% for NTL9; the average pairwise identity between consensus sequences and each sequence in the corresponding alignment ranges from 58% for NTL9 and PGK to 40% for SH3 and SH2 (SI Appendix, Figure S2).

### Structural characterization of consensus proteins

To determine whether the six consensus sequence proteins adopt their target folds, we expressed and purified consensus proteins (hereafter denoted with a c, e.g., cNTL9) for each of the six protein families and characterized them using CD and NMR spectroscopies. Far-UV CD spectra for cNTL9, cAK, and cPGK show minima at 222 and/or 208 nm (Figure 2A), consistent with α-helical secondary structure. Spectra for cSH2 and cDHFR show single minima at 219 and 214 nm respectively, consistent with predominantly β-sheet secondary structure. Consensus SH3 shows a far-UV CD spectrum with minima at 228 and 204 nm and a maximum at 219 nm, similar to published spectra for a naturally-occurring SH3 domain (37).

**Figure 2.**
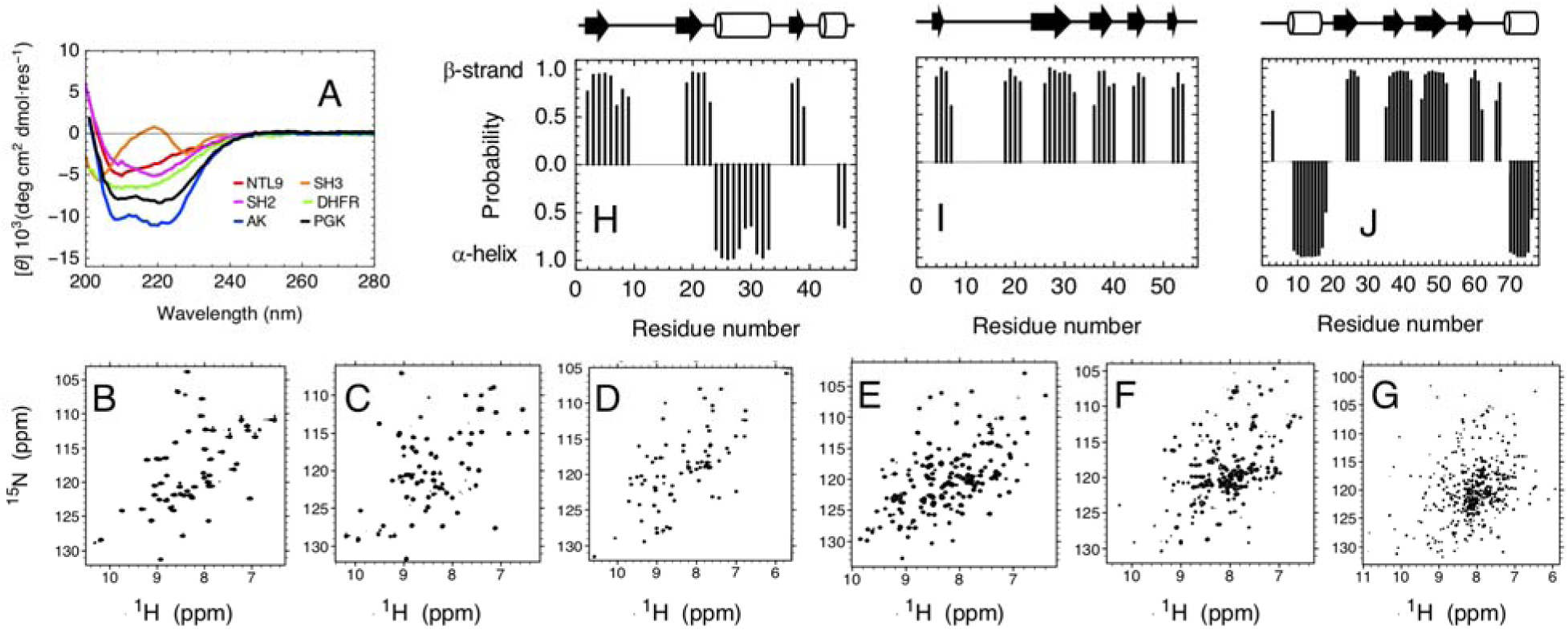
Structural features of consensus proteins. (A) Far-UV CD spectra for all consensus proteins. (B-F) ^1^H-^15^N HSQC spectra for (B) cNTL9, (C) cSH3, (D) cSH2, (E) cDHFR with addition of 1:1 molar equivalents of methotrexate, and (F) cAK at 600 MHz. (G) ^1^H-^15^N TROSY spectrum for consensus cPGK at 800 MHz. (H-J) Secondary chemical shift-based secondary structure probabilities using backbone resonances calculated by TALOS-N for (H) cNTL9, (I) cSH3, and (J) cSH2. Schematics above plots represent sequence positions of secondary structures in homology models of consensus sequences.

With the exception of cDHFR, ^1^H-^15^N HSQC and TROSY (for cPGK) NMR spectra for each consensus protein show sharp resonances that are well dispersed in the ^1^H dimension (Figure 2B--G), indicating that these consensus proteins adopt well-folded tertiary structures. These six consensus proteins show between 88--99% of expected cross-peaks in the ^1^H-^15^N HSQC and TROSY (cPGK) spectra, suggesting that the consensus proteins maintain rigid tertiary structure over most of their sequence. The high signal-to-noise and sharp cross-peaks indicate that these consensus proteins have high solubility at NMR concentrations (ranging from 400--800 μM).

For cDHFR, the ^1^H-^15^N HSQC spectrum shows rather broad resonances and only around 60% of expected cross peaks (SI Appendix, Figure S3). However, addition of the substrate analog methotrexate both sharpens resonances and increases the number of resolved cross peaks (Figure 2E), suggesting that cDHFR may undergo a large-scale conformational change and rigidification upon binding. This behavior has been observed in a study of two naturally-occurring DHFR proteins (38).

To determine the location of α-helices and β-strands, we assigned backbone resonances for cNTL9, cSH3, and cSH2. Using standard triple-resonance experiments, we were able to assign backbone ^1^H and ^15^N resonances, as well as ^13^C Cα, Cβ, and C’ chemical shifts, for 96%, 96%, and 89% of residues for cNTL9, cSH3, and cSH2 respectively (SI Appendix, Figures S4--S6). Secondary chemical shift-based secondary structure predictions using these resonances show α-helix and β-strand boundaries that largely match secondary structure locations in consensus homology models, suggesting that the consensus proteins adopt their archetypal folds (Figure 2H--J).

### Equilibrium stability of consensus proteins

To determine equilibrium folding free energies, we measured guanidine hydrochloride-(GdnHCl) and temperature-induced unfolding transitions for all consensus proteins using a combination of CD and fluorescence spectroscopies. All consensus proteins show sigmoidal GdnHCl-induced unfolding transitions (Figure 3A, SI Appendix, Figure S8), indicating that the consensus proteins unfold in a cooperative manner similar to naturally-occurring proteins. Four of the six consensus proteins showed sigmoidal thermal unfolding transitions, with T_m_ values ranging from 62 °C to 87°C. For cNTL9 and cSH2, unfolding transitions were not observed up to the maximal temperature of 93 °C; addition of 4 and 2 M GdnHCl revealed thermal unfolding transitions, demonstrating that these proteins are hyperstable, with thermal unfolding transitions significantly above 93 °C in the absence of denaturant.

**Figure 3.**
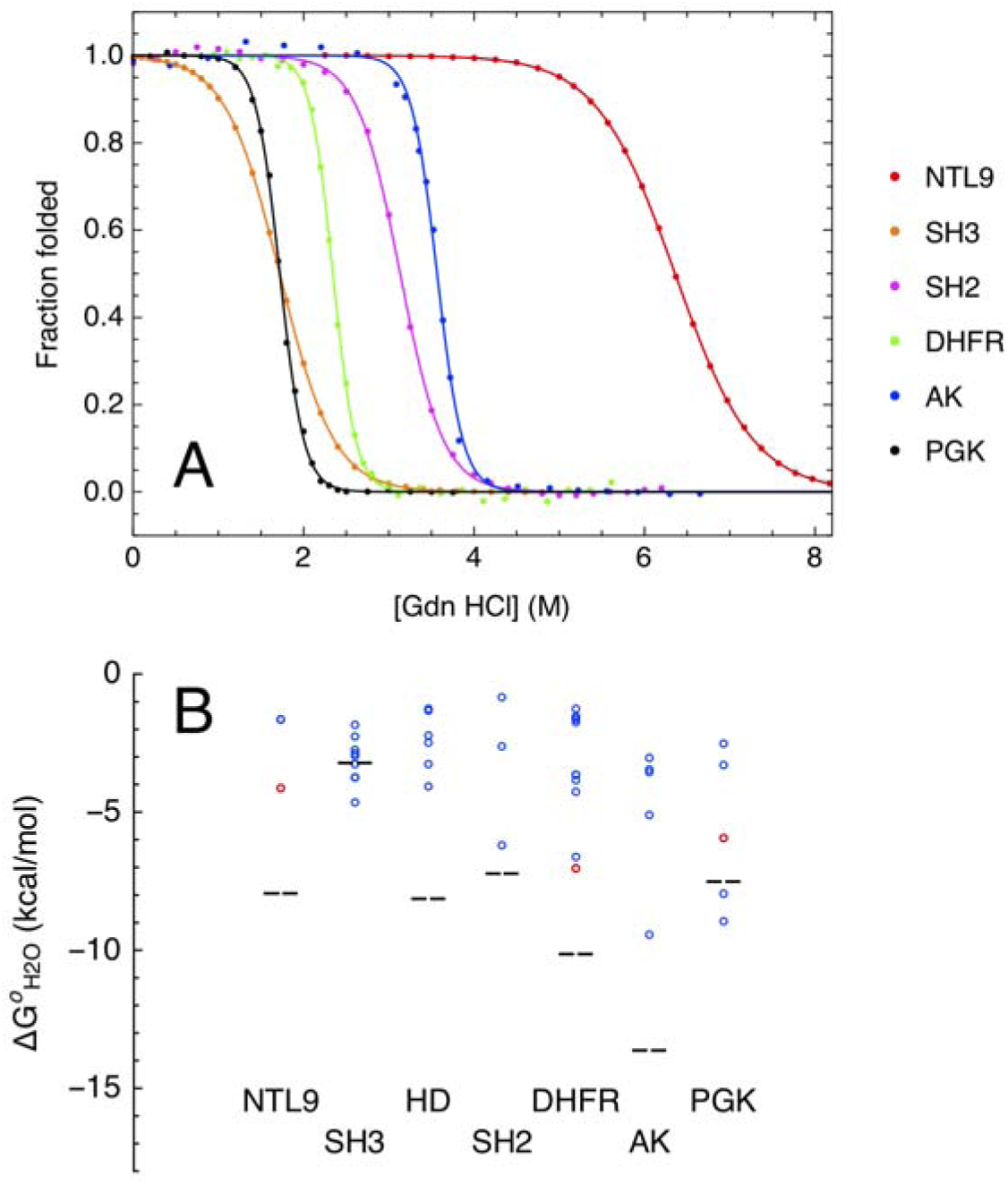
Equilibrium folding of consensus proteins. (A) Representative guanidine hydrochloride-induced folding/unfolding curves for consensus proteins. Data were collected using circular dichroism (cDHFR, cAK, cPGK) or fluorescence (cNTL9, cSH3, cSH2) spectroscopies. Solid lines are obtained from fitting a two-state model to the data. (B) Comparison of measured folding free energies of consensus proteins to those of extant sequences. Open circles represent folding free energies of extant eukaryotic or mesophilic (blue) and thermophilic (red) sequences. Black dashed lines represent measured folding free energies of consensus sequences. Folding free energy values and sources are reported in SI Appendix, TableS3.

Of the four proteins that showed thermal unfolding transitions in the absence of GdnHCl, two (cSH3 and cAK) were not fully reversible. In contrast, by chemical denaturation, all consensus proteins were found to fold and unfold reversibly, allowing us to determine equilibrium folding free energies and *m*-values by fitting a two-state linear extrapolation model to all consensus protein GdnHCl denaturations. Folding free energies ranged from −3.2 kcal/mol for cSH3 to −13.6 kcal/mol for cAK (Table 1). Measured *m*-values for cNTL9 and cSH2 match values predicted from an empirical relationship based on chain-length (39, 40) within 5%, suggestive of two-state folding. The measured *m*-value for cSH3 is 30% larger than the predicted value. Measured *m-value*s for cDHFR, cAK, and cPGK are 20%, 50%, and 70% smaller than predicted values, suggesting the population of partly folded states in the folding transition region. As a result, estimated folding free energies for these constructs are likely to underestimate the free energy difference between the native and denatured states.

**Table 1.** Equilibrium folding free energies of consensus proteins.

| | $\Delta G^{\circ}_{H_2O}$<br>(kcal mol<sup>-1</sup>) | $m$-value<br>(kcal mol<sup>-1</sup><br>M<sup>-1</sup>) | $\Delta\Delta G^{\circ}_{H_2O}$<br>(kcal mol<sup>-1</sup>) |
| --- | --- | --- | --- |
| cNTL9 | -7.9 ± 0.1 | 1.23 ± 0.02 | -4.7 |
| cSH3 | -3.2 ± 0.1 | 1.88 ± 0.05 | +0.3 |
| cHD | -10.8 ± 0.5 | 1.60 ± 0.08 | -5.3 |
| cSH2 | -7.2 ± 0.1 | 2.28 ± 0.01 | -3.6 |
| cDHFR | -10.1 ± 0.1 | 4.32 ± 0.03 | -6.7 |
| cAK | -13.6 ± 0.5 | 3.83 ± 0.13 | -8.3 |
| cPGK | -7.5 ± 0.2 | 4.34 ± 0.12 | -1.4 |

To compare the measured folding free energies of the consensus proteins with extant sequences, we gathered folding free energy values from the literature for naturally-occurring sequences for each protein family. We limited our search to folding free energy values determined by chemical denaturation experiments and to proteins that appear to be monomeric^‡^. For comparison, we have included the consensus homeodomain (cHD) that we recently characterized with this approach (32).

The stabilities of six of the seven consensus proteins are greater than the mean stabilities of the extant homologues (Figure 3B; SI Appendix, Table S3), with stability increases ranging from 8.3 kcal/mol for consensus AK to 1.4 kcal/mol for cPGK (although for PGK, fitted ΔG° parameters are likely to underestimate the free energy difference between native and denatured ensembles since the unfolding transition is almost certainly multistate). The sole exception to this observation is cSH3, which shows a stability that is 0.3 kcal/mol less than the mean value for the extant homologues. Furthermore, stabilities of five of the seven consensus proteins (cNTL9, cHD, cSH2, cDHFR, and cAK) are greater than that of the most stable extant homologue, ranging from 0.6 kcal/mol for cSH2 to 3.8 kcal/mol for cAK (Figure 3B; SI Appendix, Table S3).

### Characterization of cSH3 peptide binding

To measure binding of cSH3 to a target peptide, we acquired a synthetic peptide (Ac-PLPPLPRRALSVW-NH_2_), which contains a proline-rich motif used in previous binding studies of the human Fyn SH3 domain, a well-studied extant sequence with peptide contact residues that match those in consensus SH3 (41, 42). ^1^H-^15^N HSQC spectra of ^15^N-labelled cSH3 at increasing concentrations of unlabeled peptide show shifts in some (but not all) peaks (SI Appendix, Figure S9). This behavior is consistent with formation of a complex that is in fast exchange on the chemical shift timescale. When plotted on a homology model of cSH3, the largest chemical shift perturbations (CSPs) cluster on the canonical peptide-binding site (Figure 4A). A global fit of a single-site binding model to the 10 residues displaying the largest CSPs gives a K_d_ of 795 ± 13 μM (Figure 4B; the reported uncertainty is at a 67% confidence level derived from 5000 bootstrap iterations in which residuals are randomly resampled with replacement). This measured binding affinity for cSH3 is approximately 5000-fold weaker than a binding affinity reported for human Fyn SH3 domain binding the same peptide sequence (41).

**Figure 4.**
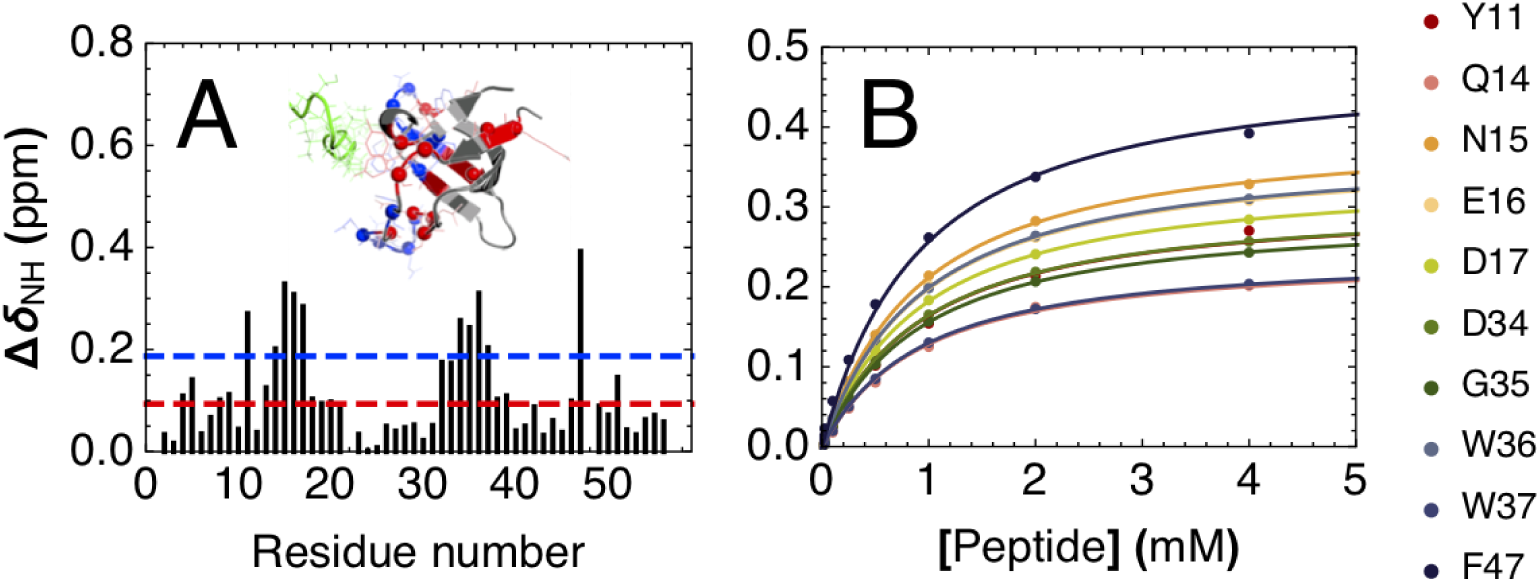
Peptide binding of cSH3 by NMR spectroscopy. (A) Chemical shift perturbations (CSPs) of backbone ^1^H-^15^N HSQC cross-peaks resulting from addition of a 20-fold excess unlabelled peptide to ^15^N-labelled consensus SH3. Red and blue lines indicate one and two standard deviations respectively. Inset shows a homology model of cSH3 aligned to a peptide-bound structure of human Fyn SH3 (PDB 1A0N; peptide shown in green). Residues showing CSPs greater than one and two standard deviations are shown with alpha carbons as spheres, side chains atoms as lines, and colored red and blue respectively. (B) Binding isotherms for the ten consensus SH3 residues showing largest CSPs. Solid lines are obtained from a global fit to a single-site binding model using a common dissociation constant for all residues.

### Steady-state kinetics of consensus enzymes

To investigate how the consensus design strategy affects enzymatic activity, we characterized steady-state kinetics of catalysis for the three consensus enzymes in our set. DHFR, AK, and PGK catalyze various chemical reactions important for cellular metabolism:

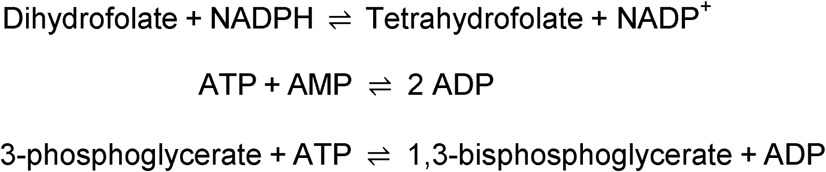

For each enzyme, we measured reaction velocities at 20 °C as a function of substrate concentration, monitoring the oxidation of NAD(P)H spectrophotometrically either as a direct read-out of activity (DHFR) or in an enzyme-coupled reaction (see Materials and Methods). Strikingly, all three consensus enzymes show substantial activity (Figure 5). Michaelis constants for the consensus enzymes are similar to those of extant enzymes, ranging from 10-fold higher to 3-fold lower than values determined under similar conditions (Table 2). Steady-state k_cat_ values determined at 20 °C for the consensus enzymes are all smaller than those for mesophilic homologues (Table 2), but range from comparable to (1.3-fold lower) to slightly lower (roughly 6-fold lower) k_cat_ values for thermophilic homologues.

**Table 2.**
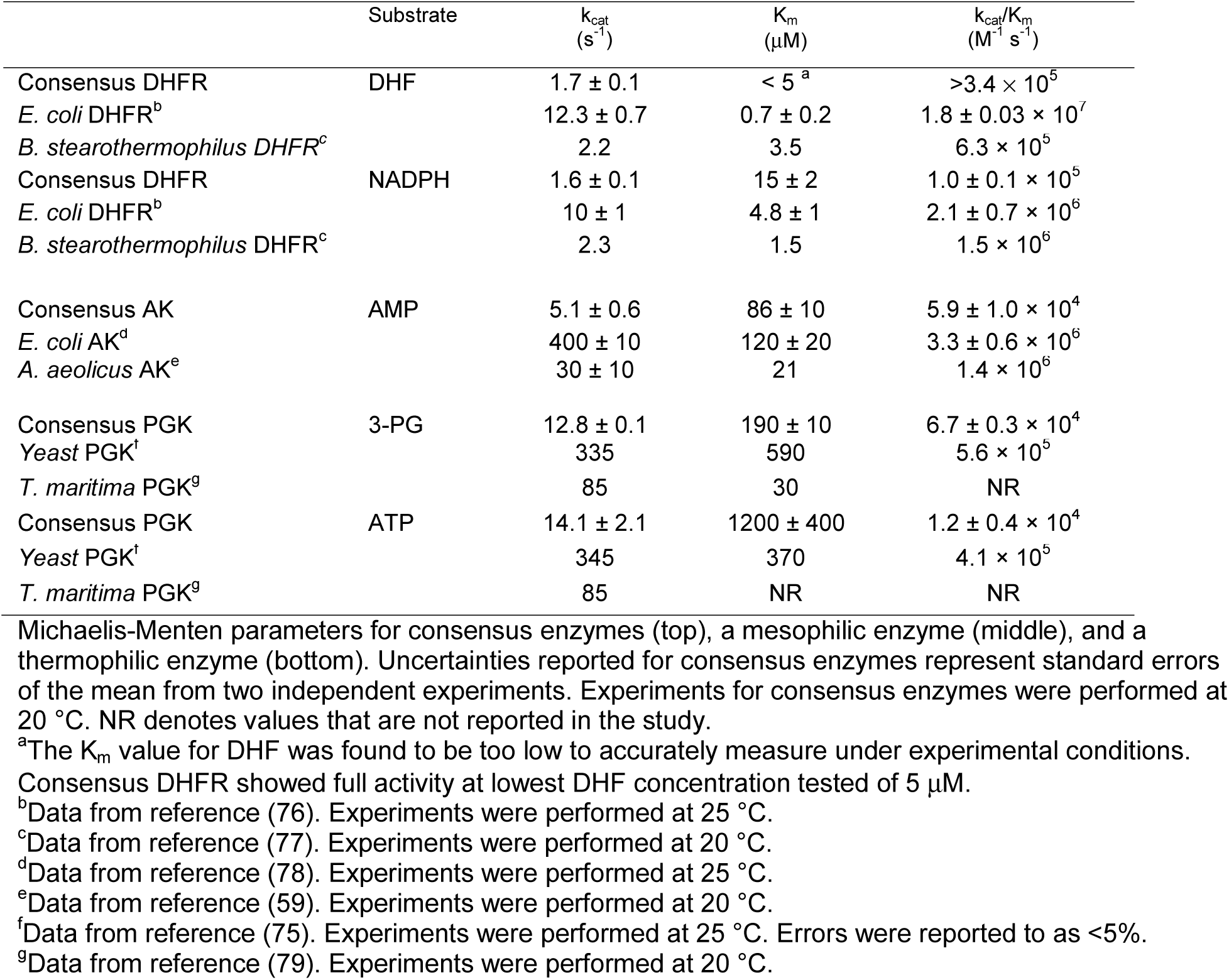
Steady-state kinetic parameters of catalysis for consensus and naturally-occurring enzymes.

| | Substrate | $k_{cat}$<br>(s<sup>-1</sup>) | $K_m$<br>(μM) | $k_{cat}/K_m$<br>(M<sup>-1</sup> s<sup>-1</sup>) |
| --- | --- | --- | --- | --- |
| Consensus DHFR | DHF | 1.7 ± 0.1 | < 5<sup>a</sup> | >3.4 $\times 10^5$ |
| *E. coli* DHFR<sup>b</sup> | | 12.3 ± 0.7 | 0.7 ± 0.2 | 1.8 ± 0.03 $\times 10^7$ |
| *B. stearothermophilus* DHFR<sup>c</sup> | | 2.2 | 3.5 | 6.3 $\times 10^5$ |
| Consensus DHFR | NADPH | 1.6 ± 0.1 | 15 ± 2 | 1.0 ± 0.1 $\times 10^5$ |
| *E. coli* DHFR<sup>b</sup> | | 10 ± 1 | 4.8 ± 1 | 2.1 ± 0.7 $\times 10^6$ |
| *B. stearothermophilus* DHFR<sup>c</sup> | | 2.3 | 1.5 | 1.5 $\times 10^6$ |
| Consensus AK | AMP | 5.1 ± 0.6 | 86 ± 10 | 5.9 ± 1.0 $\times 10^4$ |
| *E. coli* AK<sup>d</sup> | | 400 ± 10 | 120 ± 20 | 3.3 ± 0.6 $\times 10^6$ |
| *A. aeolicus* AK<sup>e</sup> | | 30 ± 10 | 21 | 1.4 $\times 10^6$ |
| Consensus PGK | 3-PG | 12.8 ± 0.1 | 190 ± 10 | 6.7 ± 0.3 $\times 10^4$ |
| Yeast PGK<sup>f</sup> | | 335 | 590 | 5.6 $\times 10^5$ |
| *T. maritima* PGK<sup>g</sup> |  | 85 | 30 | NR |
| Consensus PGK | ATP | 14.1 ± 2.1 | 1200 ± 400 | 1.2 ± 0.4 $\times 10^4$ |
| Yeast PGK<sup>f</sup> | | 345 | 370 | 4.1 $\times 10^5$ |
| *T. maritima* PGK<sup>g</sup> |  | 85 | NR | NR |

Michaelis-Menten parameters for consensus enzymes (top), a mesophilic enzyme (middle), and a thermophilic enzyme (bottom). Uncertainties reported for consensus enzymes represent standard errors of the mean from two independent experiments. Experiments for consensus enzymes were performed at 20 °C. NR denotes values that are not reported in the study.
<sup>a</sup>The $K_m$ value for DHF was found to be too low to accurately measure under experimental conditions.
Consensus DHFR showed full activity at lowest DHF concentration tested of 5 μM.
<sup>b</sup>Data from reference (76). Experiments were performed at 25 °C.
<sup>c</sup>Data from reference (77). Experiments were performed at 20 °C.
<sup>d</sup>Data from reference (78). Experiments were performed at 25 °C.
<sup>e</sup>Data from reference (59). Experiments were performed at 20 °C.
<sup>f</sup>Data from reference (75). Experiments were performed at 25 °C. Errors were reported to as <5%.
<sup>g</sup>Data from reference (79). Experiments were performed at 20 °C.

**Figure 5.**
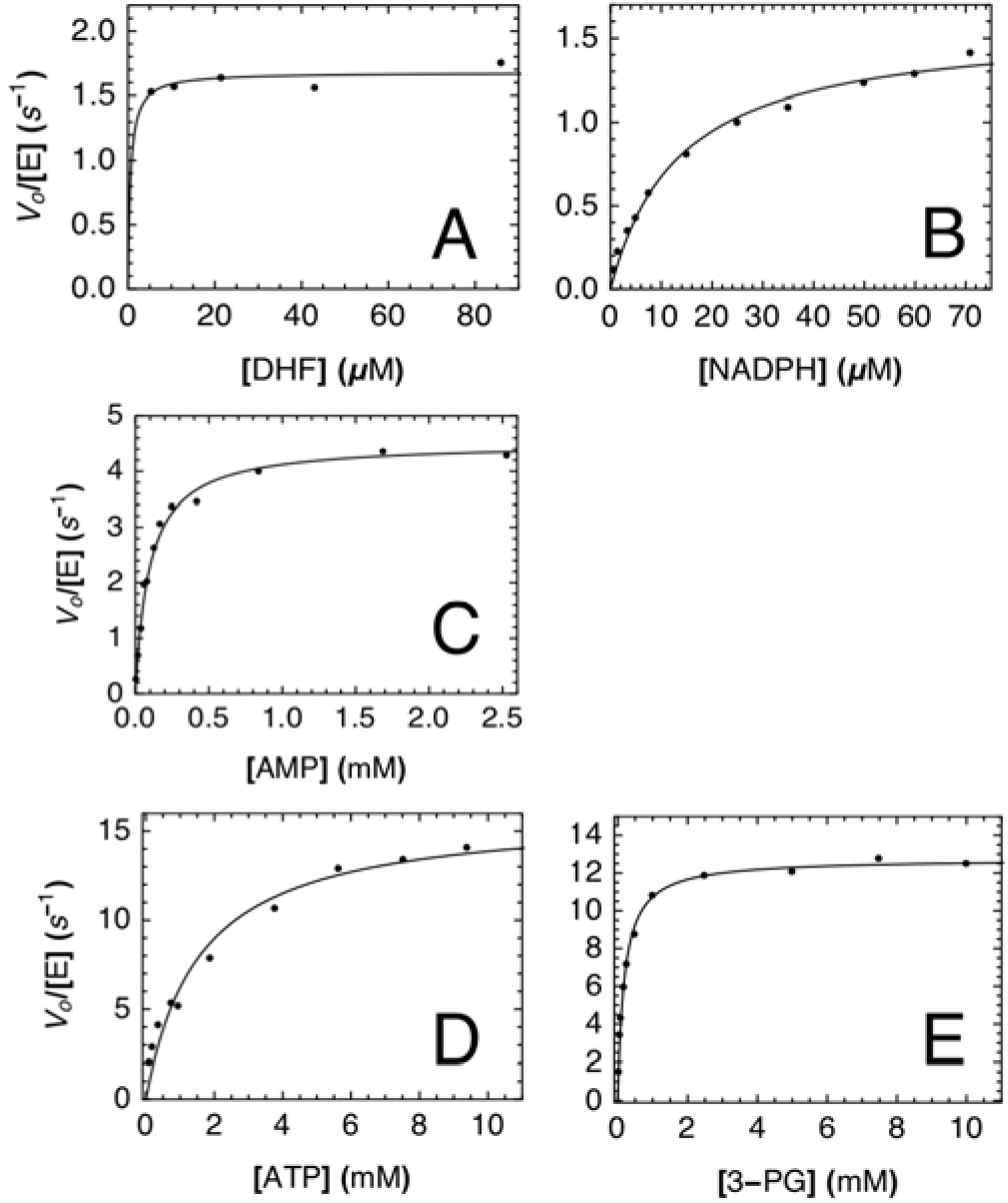
Steady-state kinetics of consensus enzymes. Representative Michaelis-Menten curves generated for each substrate for cDHFR (A, varying DHF with 150 μM NADPH; B, varying NADPH with 100 μM DHF), cAK (C, varying AMP with 5 mM ATP), and cPGK (D, varying 3-PG with 20 mM ATP; E, varying ATP with 10 mM 3-PG). Solid lines are obtained from fitting a Michaelis-Menten model to the data. In (A), full activity was obtained at the lowest DHF concentration we could measure; as a result, the K_m_ value for DHF could not be accurately determined.

The high thermal stability of the consensus enzymes allows us to measure activity at high temperatures where reaction rates likely increase considerably, providing a better comparison to thermophilic homologues. To determine the effects of temperature on enzyme activity, we measured k_cat_ values for the consensus enzymes over a range of temperatures. For cDHFR, where absorbance provides a direct readout of catalysis, we were able to access temperatures up to 50 °C. However, for cPGK and cAK, we were restricted to 40 °C due to the limited stability of the coupling enzymes used in the absorbance assay. For cAK, we were able to expand this temperature range to 70 °C using a ^31^P NMR-based assay (AK; see SI Appendix and reference (43), which directly monitors conversion of reactants to products. Over the accessible temperature range, all consensus enzymes show exponential increases in catalytic rates with increased temperature, consistent with Arrhenius kinetics (SI Appendix, Figure S10), allowing us to extrapolate k_cat_ values to other temperatures. On the whole, the consensus enzymes show levels of activity comparable to thermophilic homologues (SI Appendix, Table S4), although there are variations depending on the consensus enzyme and the thermophile being compared. Consensus DHFR has a larger turnover number than two thermophilic homologues (by 5- and 31-fold). Consensus PGK shows both larger (1.8-fold increase) and smaller turnover numbers (1.2-, 4-, 5-, and 13-fold decreases). In contrast, cAK has a smaller turnover number (by 21-fold).

### Sequence properties of consensus proteins and consensus mismatches

The studies above show that for all six families examined here, consensus design is successful. That is, proteins adopt well folded structures, these structures match the fold of the families from which they derived, they retain biological activity, and importantly, their stabilities equal or (more often) exceed median stabilities of extant. With this broad collection of consensus-designed proteins, we are in the position to ask whether there are any sequence features that set these consensus proteins apart from extant sequences, and if so, which features are most important for increased stability. Such properties include general position-independent features such as sequence composition, charge, polarity, and hydrophobicity, as well as position-specific properties within each family such as degree of conservation and surface accessibility.

#### General sequence features

We compared various sequence properties our six consensus sequences to those of the naturally-occurring sequences within our curated multiple sequence alignments. As with the stability comparisons above, we have included a consensus homeodomain that we have described previously (32). For every sequence (consensus and every sequence in each MSA), we calculated the proportion of sequence (averaging over all sites) made up of charged residues (D, E, K, R), polar uncharged residues (C, H, N, Q, S, T), total polar residues (the sum of charged and polar uncharged), and nonpolar residues. Similarly, we calculated the net charge of each sequence (the difference between the number of positively and negatively charged residues, assuming full positive charges for R and K, and full negative charges for D and E).

For each metric, the naturally-occurring sequences appeared approximately normally distributed (SI Appendix, Figure S11). Consensus sequences show strong biases within the distributions of natural sequences for some of these sequence and structural features. Most notably, consensus sequences all lie towards the upper-tail end of the distributions for proportion of charged residues, averaging 2.1 standard deviations above the mean (Figure 6A). Conversely, consensus sequences lie toward the lower-tail end of the distributions for proportion of polar uncharged residues, averaging 2.1 standard deviations below the mean. These two biases offset in terms of total polar (charged and uncharged) residues: consensus sequences lie near the middle of the distribution for the proportion of total polar residues, averaging 0.1 standard deviations below the mean. Likewise, consensus sequences lie near the middle of the distribution for the proportion of nonpolar residues, as would be expected given the proportion of total polar residues. Thus, it appears that the consensus design strategy implicitly creates sequences that are preferentially enhanced in charged residues at the expense of polar uncharged residues, without a large perturbation of the number of total polar versus nonpolar residues. Two protein families, SH3 domain and homeodomain, deviate from this trend, both showing and an increase in total polar residue content and a decrease in nonpolar residue content. The net charge of six of the seven consensus protein sequences are also close to mean values from the MSA, with DHFR showing the only significant deviation.

**Figure 6.**
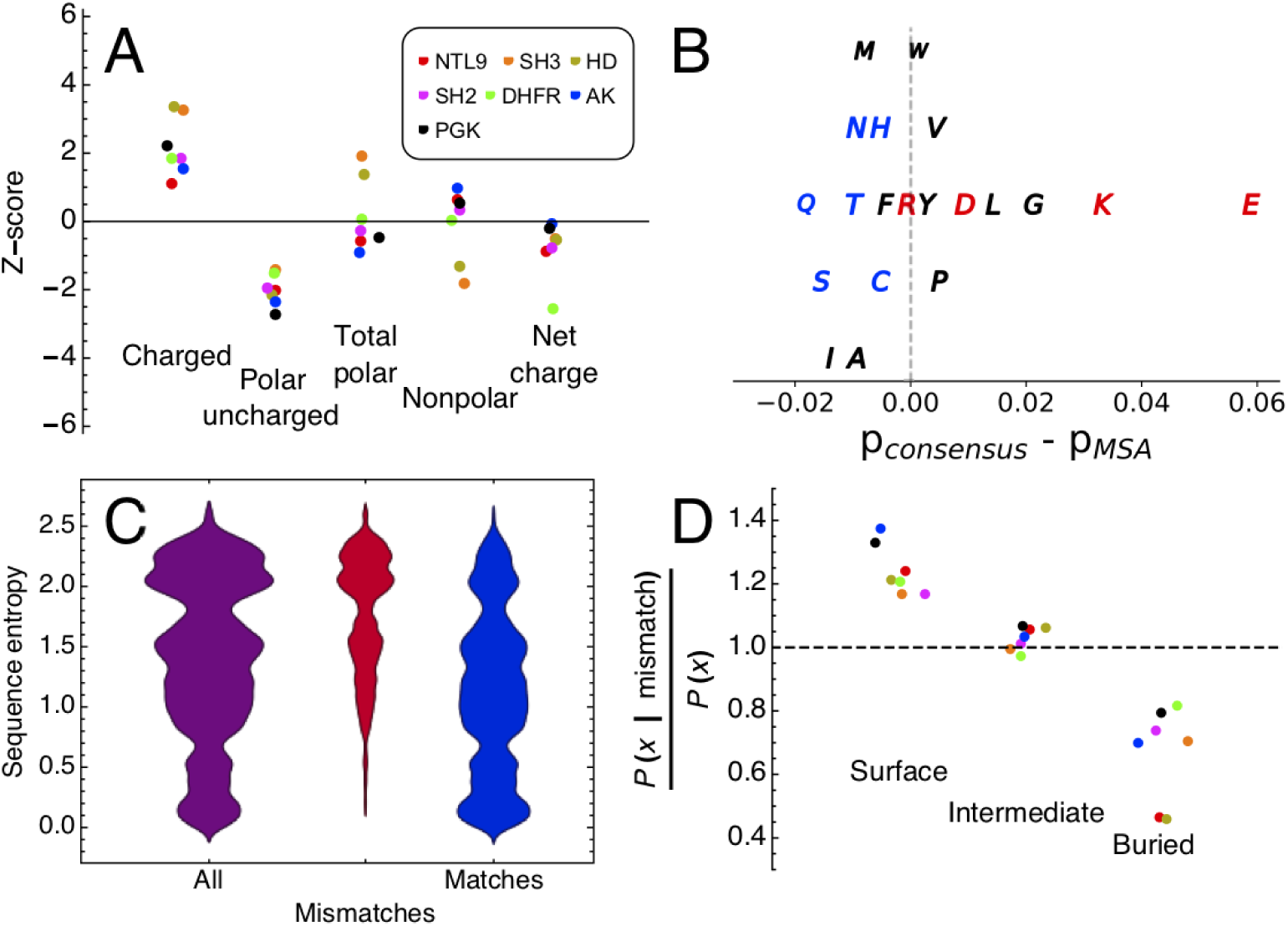
Sequence properties of consensus sequences. (A) Z-scores (the number of standard deviations that separate the consensus sequence from the mean value of sequences in the multiple sequence alignments, Equation 3 in SI Appendix) for various sequence properties. Distributions for all protein families are shown in SI Appendix, Figure S11. (B) Differences between residue frequencies in the consensus sequences and the MSAs averaged over all seven protein families. Residues are colored as: polar charged (red), polar uncharged (blue), nonpolar (black). The vertical offset is used for clarity. (C) Distributions of sequence entropy values for all positions in the PGK MSA (purple), positions at which residues in extant sequences differ from the consensus sequence (consensus mismatches; red), and positions at which residues in extant sequences match the consensus sequence (consensus matches; blue) for PGK. Sequence entropy distributions for all protein families are shown in SI Appendix, Figure S13. (D) Ratios of conditional probabilities of different structural environments (surface, intermediate, and buried; "X" in the y-label) for consensus mismatches relative to overall probabilities of surface, intermediate, and buried residues at all positions. Conditional and overall probabilities for all protein families are shown in SI Appendix, Figure S14. Legend as in panel A.

The observed increase in the proportion of charged residues in consensus sequences relative to extant sequences must arise from differences in the proportions of each of the charged residues individually. However, an increase in the charged residue content does not require that the proportion of each charged residue increases. To determine the specific residues responsible for this overall increase, we compared the proportions of individual residues over the extant sequences in each MSA (again averaging over all sites) to those in the consensus sequence. Many residues show consistent enrichment or depletion in consensus sequences relative to the extant sequences. For example, consensus sequences are enriched in of D, E, and K and depleted in C, N, Q, S, and T in all or six of the seven protein families (SI Appendix, Figure S12). These trends are highlighted when the relative differences between the consensus sequence and extant sequences for each residue are averaged over the seven protein families. Consensus sequences show the largest enrichments in E, K, G, L, and D, the largest depletion in Q, S, I, and marginal effects on all other residues (Figure 6B). Thus, the increase in charged residues for the consensus proteins results from increased proportions of D, E, and K, but not R, while the decrease in polar uncharged residues results from decreased proportions (albeit modest proportions for some residues) of all polar uncharged residues.

#### Position-specific sequence features

In addition to the general sequence features described above, which average over each sequence, we also examined the features of sequence substitution in a position-specific way. At each position in the MSA, we compared the residue of each extant sequence to the consensus residue at that position. We separated these residue-specific comparisons into “consensus matches” and “consensus mismatches” for the positions at which extant residue matched or differed from the consensus residue, respectively. Clearly, all changes in structure, stability, and function in consensus constructs compared to extant homologues result from the residues that differ from the consensus. Thus, the properties of these "consensus mismatches" are important for understanding, for example, the increase in stability we have observed for most of our consensus constructs.

On average, there will be more consensus mismatches at positions with low sequence conservation, although there will also be some number of mismatches at positions with high conservation. One simple question that can be asked about consensus mismatches is "how biased are consensus mismatches towards positions of low conservation?" To quantify the extent to which consensus mismatches are made at sites with high conservation, we calculated the sequence entropy (44) at each position *i* in each MSA using the formula

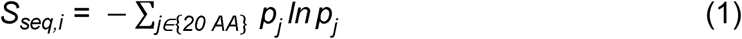

where *p*_*j*_ is the frequency of residue *j* occurring at position *i*, and plotted this distribution for each family (purple distributions in Figure 6C; SI Appendix, Figure S13). We then weighted this distribution by the fraction of sequences in the multiple sequence alignment that differ from the consensus at each position

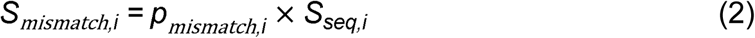

(red distributions in Figure 6C; SI Appendix, Figure S13) and the fraction that match the consensus at each position

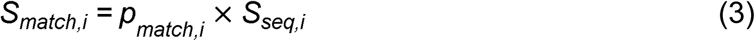

(blue distributions in Figure 6C; SI Appendix, Figure S13).

As expected, we see a bias of consensus matches towards low sequence entropy positions and a bias of consensus substitutions towards high entropy positions. For example, at positions that have entropies below 0.5, nearly all sequences in the MSA match the consensus. Likewise, at positions that have entropies above value of 2, most sequences in the MSA differ from consensus. However, at positions with intermediate entropies (ranging from 1 to 1.5), there are significant numbers of sequences in each MSA that differ from consensus. It is possible that these substitutions disproportionately contribute to the observed stability increase of consensus proteins.

Finally, we examined the extent to which these sequence differences occur at surface positions. Since surface residues tend to contribute less to overall stability than buried residues (45, 46), we might expect the stability increases of consensus proteins to have their origins in substitutions at buried positions. However, because surface positions tend to be more highly variable, we expect most consensus substitutions to occur at surface sites; as a result, surface residues may contribute to stability enhancement through the collective effect from many substitutions. Consistent with this, Makhatadze and coworkers have shown that stability can be increased through careful placement of surface charged residues (47, 48). To identify surface positions within each multiple sequence alignment, we generated a homology model from the corresponding consensus sequence, calculated the side chain solvent accessible surface area at each position, and referenced to the average side chain solvent accessible surface area of the same residue in an extended peptide (see Materials and Methods in SI Appendix).

A comparison of the distribution of consensus mismatches occurring at surface, intermediate, and buried positions shows that for six of the seven proteins, consensus mismatches occur most frequently at surface positions (SI Appendix, Figure S14; darker bars). Overall, mismatches occur at surface, intermediate, and buried positions with median proportion of 0.49, 0.27, and 0.22 respectively. However, these proportions are dependent on the overall proportions of surface, intermediate, and buried residues that make up each protein. The four smaller domain families of NTL9, SH3, HD, and SH2 all have more surface residues than intermediate or buried residues (SI Appendix, Figure S14; lighter bars). However, the larger proteins DHFR, AK, and PGK have more buried residues than surface or intermediate residues (consistent with a decreased surface to volume ratio for larger proteins).

To account for this underlying difference in surface accessibility, we divided the conditional probabilities that consensus mismatches are located at surface, intermediate, and buried positions (i.e., p(X | mismatch), where X represents the three structural classes; dark bars in SI Appendix, Figure S14) by the overall marginal probabilities of the three structural classes (surface, intermediate, buried; light bars in SI Appendix, Figure S14). By Bayes’ theorem, this ratio represents the true enrichment (or supression) of consensus mismatches at surface, intermediate, and exposed sites. This ratio shows an enrichment of consensus mismatches at surface positions for all seven protein families (Figure 6D), with an average factor of 1.21. In contrast, there is a suppression of consensus mismatches at core positions in all seven protein families, with an average factor of 0.71. At intermediate positions, mismatches are neither enhanced nor supressed (Figure 6D, with an average factor of 1.03).

## Discussion

There are a number of reports in the literature in which consensus design has successfully been used to produce proteins that adopt their archetypical fold. This approach has found recent success for linear repeat-protein targets (49–52), but has also shown some success for globular protein targets (13, 14, 23, 26, 27). In most reports, globular consensus proteins have enhanced equilibrium stability, and sometimes retain biological activity. However, point-substitution studies suggest stability gains from wholesale consensus substitution may be marginal, and protein engineering studies with a WW domain suggest that consensus proteins may lack the coupling energies needed to fold (53). Because publications of successful design of globular proteins have so far been “one target at a time”, and since unsuccessful designs are unlikely to be published, it is difficult to estimate how likely the consensus approach is to yield folded proteins, and to determine how to improve on the approach when it performs poorly. The studies presented here, where we generate and characterize consensus sequences for six protein families, provide an unbiased look at rates of success and failure. Comparisons for different targets are aided by the use of a common design protocol for all targets, a common purification pipeline, and a uniform set of protocols for measuring stability, structure, and function. This multi-target approach also allows us to evaluate sequence features that are likely to contribute consensus stabilization in general way, and correlate these features with measured biochemical properties.

Our results show that consensus sequence design is both a general and successful strategy for small and large globular proteins with diverse folds. For all six protein families targeted, the resulting consensus proteins expressed to high levels, remained soluble in solution, and adopted a well-folded tertiary structure. Structural information obtained from CD and NMR spectroscopies suggest that all of the consensus proteins adopt their archetypal fold. Support for folding is provided by the observation that all consensus proteins tested show their expected biological activities (Figures 2, 4, and 5).

Our findings indicate that thermodynamic stabilization of consensus proteins is the norm, rather than the exception. Four of the six (and five of the seven including the previously reported homeodomain (32)) consensus proteins showed stabilities greater than the most stable naturally-occurring sequences identified in our literature search, and the other two showed stabilities near and above average (Figure 3; SI Appendix, Table S3). In short, consensus design can be expected to provide stability enhancements about three quarters of the time. This result is particularly surprising, given that proteins are not under direct evolutionary selection for maximal stability (54, 55), but must simply retain sufficient stability to remain folded. It appears that by taking the most probable residue at each position, a large number of small stability enhancements sum to a large net increase in stability. Although naturally occurring extant proteins are of lower stability (perhaps because high stabilities are not required for function) this information gets encoded in alignments of large numbers of sequences. Taken at face value, these results highlight the importance of information encoded at the level of single residues. Though single-residue information does not explicitly include pairwise sequence correlations, favorable pairs may be retained in our consensus designs as long as each residue in the pair is highly conserved. Thus, the extent to which consensus sequences capture such “accidental” correlations and their contributions to the observed effects on stability require further studies.

The source of the relative instability of cSH3 is unclear. The consensus SH3 and the extant sequences from which it was generated have some exceptional features, although none of these are unique to SH3. The SH3 sequences are among the shortest sequence families examined (54 residues) although not the shortest (NTL9 is 46 residues). The consensus SH3 sequences has a high fraction of polar residues and the lowest fraction of uncharged residues (Figure 6A), although the consensus homeodomain is a close second. The SH3 sequence is also the only protein in our set that has an all-β structure.

Taxonomically, sequences in the SH3 MSA show the lowest pairwise identity (26%; SI Appendix, Table S1), although the SH2 family is a close second (28%). It may be noteworthy that the increase in stability of cSH2 compared to the available extant values is the smallest stability increment observed for those proteins likely to conform to a two-state mechanism. It may also be noteworthy that along with SH2, the SH3 multiple sequence is uniquely dominated by eukaryotic sequences. In a study of consensus superoxide dismutase sequences, Goyal and Magliery found successful consensus design to be highly dependent on the phylogeny represented in the MSA (56).

A particular advantage of consensus sequence design is that it draws upon the natural evolutionary history of a protein family. As a result, residues important for function are likely to be conserved (57). However, it cannot be assumed *a priori* that the resulting consensus proteins will show biological function, since consensus sequences are novel sequences that have experienced no evolutionary selection for function. Importantly, all consensus proteins we assayed for function maintained some level of expected biological activities of both molecular recognition and enzymatic catalysis (Figures 4 and 5). This result, combined with previously reported studies showing consensus protein function (22–31), indicates that information necessary for protein function is retained in averaging over many sequences that each individually contain functionally important information.

In both this study and our previous investigations of a consensus homeodomain (32), consensus substitution showed varying effects on molecular recognition. Consensus HD and cSH3 each showed 2-3 orders of magnitude differences in their binding affinities to cognate substrates relative to naturally-occurring sequences, with cHD showing higher affinity and cSH3 showing lower affinity. The origins of these differences in substrate binding affinities remain unclear. It is possible that the sequences used to obtain the consensus homeodomain sequence bind similar sequences (indeed, many of these sequences are from the engrailed superfamily), resulting in an “optimized” homeodomain, whereas sequences used to obtain the consensus SH3 possess different specificities, resulting in a sequence whose binding affinity has been “averaged out.” Testing this explanation will require an investigation of the binding specificities of the consensus proteins as well as those of the sequences used to generate them.

For the three enzymes examined here, consensus substitution shows variable effects on catalysis. At low temperatures (20 °C), steady-state turnover numbers for all consensus enzymes were smaller than those for naturally-occurring mesophilic sequences, but on par with those of thermophilic homologues (Table 2). This is consistent with the observation that thermophilic proteins show lower catalytic activities at low temperatures than their less stable mesophilic counterparts (58, 59). At higher temperatures, cPGK has a k_cat_ value comparable to thermophilic homologues, whereas cDHFR and cAK have higher and lower k_cat_ values, respectively. On the whole, this observation demonstrates that consensus enzymes can (but sometimes don’t) achieve the same level of activities as their naturally occurring counterparts. The observed inverse correlations between enzyme stability and catalytic rates have widely been interpreted as resulting from a trade-off between dynamics and catalysis (60). As this issue is still debated (61, 62), consensus sequence design may offer a promising avenue to gain insights into the relationships among protein stability, dynamics, activity, and evolution.

The consensus design strategy used here appears to impart a strong bias on sequence: taking the most probable residue at each position does not result in average composition. This bias may have significant effects on stability and function. The consensus sequences all have a high content of charged residues and low content of polar uncharged residues (Figure 6). Consensus substitutions from uncharged to charged residues show a stronger bias towards positions of higher sequence entropy than substitutions among uncharged residues (SI Appendix, Figure S15). Thus, the overall enrichment of charged residues in consensus sequences results from charged residues (E, D, and K) “winning” over uncharged residues at positions with low conservation.

Similar (but not identical) compositional biases have also been observed in thermophilic sequences, consistent with the high stabilities observed for the consensus proteins. Like our consensus sequences, thermophilic proteins have been shown to be enriched in E and K, and depleted in A, C, H, Q, S, and T (63, 64). However, thermophilic proteins have also been shown to be enriched in Y, R, and I, which are at or below average composition in our consensus sequences. Though it might be expected that the inclusion of sequences from thermophilic organisms in our MSAs contributes both to the composition bias and to high stabilities, most of the sequences in our MSAs are from mesophilic organisms. The sequences in the SH3 and SH2 MSAs are predominantly eukaryotic (as were our previous cHD sequences); aside from a small number of moderately thermophilic fungi, these sequences all derive from mesophiles. For the other four protein families (NTL9, DHFR, AK, and PGK), MSAs are composed of at most 5% of sequences from thermophilic or hyperthermophilic bacteria or archaea (SI Appendix, Table S5). If the identified thermophilic sequences are removed from the MSA before consensus sequence generation, the resulting consensus sequences have identities of 98.6% or greater to the consensus sequence derived from the full MSAs (SI Appendix, Table S5) ^‡^.

Makhatadze et al. have been able to increase stability by introducing charged residues at surface positions of several proteins and optimizing electrostatic interactions (47, 48). It is unclear to what extent the locations of consensus charged residues optimize electrostatic interactions, or whether additional stability increases can be obtained by charge shuffling. Increases in stability and solubility have also been reported for “supercharged” proteins, which have similar numbers of charged residues (65), however, the consensus proteins studied here are generally close electroneutrality, whereas supercharged proteins have highly imbalanced positive or negative charge.

It should be noted, however, that some of the consensus sequences deviate from the general trends observed in sequence biases. For instance, consensus SH3 and HD have a greater percentage of polar residues and a lower percentage of nonpolar residues, and cDHFR has a much lower net charge, compared to the average for each MSA. Thus, consensus sequence statistics appear to abide by general trends but not absolute rules.

Similarly, analysis of the positions at which the consensus sequences differ from extant sequences highlight important aspects about the consensus design strategy. These consensus mismatches occur mainly at positions with relatively low conservation and positions on the protein surface (Figure 6), consistent with the well-known correlation between residue conservation and solvent accessible surface area (66). This may highlight an implicit advantage of consensus sequence design, since substitutions at core positions are often destabilizing (67). However, the large observed effects of consensus substitution on both stability and activity indicate that these weakly conserved and surface positions play a sizable role in both stability and function, and considerable gains can be made by optimizing these positions. This observation is consistent with the observation of the functional impacts of nonconserved "rheostat" substitutions on the surface of lac repressor (68). Furthermore, the importance of these weakly conserved positions suggests that using a large number of sequences may a key component of successful consensus design, since weakly conserved positions are most sensitive to phylogenetic noise and misalignment (69).

Our work here demonstrates that the consensus sequence design method is both a general and successful strategy to design proteins of high stability that retain biological activity. Compared to other rational, structure-based, or directed evolution methods, consensus sequence design provides a simple route to accomplish longstanding goals of protein design. Furthermore, its foundation in phylogenetics provides a promising avenue to address questions regarding the relationships of protein sequence, biophysics, and evolution.

## Materials and Methods

### Design of consensus sequences

Sequences for each family were gathered from Pfam (33), SMART (34), or InterPro (35) databases. Resulting sequence sets for each domain were filtered by sequence length, removing sequences 30% longer or shorter than the median sequence length of the set. To avoid bias from sequence groups with high identity, we used CD-HIT (70) to cluster sequences at 90% identity and selected a single representative sequence from each cluster. This curated sequence set was used to generate a multiple sequence alignment using MAFFT (71).

At each position of the multiple sequence alignment, frequencies were determined for the 20 amino acids along with a gap frequency using an in-house script. Positions occupied by residues (as opposed to a gap) in at least half of the sequences were included as positions in the consensus sequence. The most frequent residue at each of these “consensus positions” was gathered to create the consensus sequence for that protein family.

### NMR spectroscopy

^15^N- and ^13^C,^15^N-isotopically labeled proteins were expressed and purified as described in the SI Appendix. NMR samples, data acquisition, and data analysis are also described in the SI Appendix.

Peptide binding to cSH3 was monitored by heteronuclear NMR spectroscopy. A putative SH3-binding peptide (Ac-PLPPLPRRALSVW-NH_2_) was synthesized by GenScript. Samples containing 200 μM ^15^N-labeled cSH3 were prepared at 0-, 0.05-, 0.125-, 0.5-, 1.25-, 2.5-, 5-, 10-, and 20-fold molar equivalents of unlabeled peptide. ^1^H-^15^N HSQC spectra were collected on a Bruker Avance 600 MHz spectrometer in 150 mM NaCl, and 5% D_2_O, 25 mM NaPO_4_ (pH 7.0) at 25 °C. ^1^H-^15^N chemical shifts varied monotonically with peptide concentrations such that assignments from the apo-protein could be transferred to the bound state. Chemical shift perturbations were calculated and globally-fit using a single-site binding equation as described in SI Appendix.

### Circular dichroism and fluorescence spectroscopies

Circular dichroism measurements were collected on an Aviv Model 435 CD spectropolarimeter. Far-UV CD spectra were collected using a 1 mm cuvette with protein concentrations ranging from 2-31 μM at 20 °C, averaging for 5 s with a 1 nm step size. Consensus NTL9, cSH3, cSH2, and cDHFR were collected in 150 mM NaCl and 25 mM NaPO_4_ (pH 7.0). Consensus AK and cPGK were collected in 50 mM NaCl, 0.5 mM TCEP, and 25 mM Tris-HCl (pH 8.0).

GdnHCl- and temperature-induced folding/unfolding transitions were monitored using either CD or fluorescence spectroscopy. All unfolding transitions were collected with protein concentrations ranging from 1-6 μM. GdnHCl melts were collected at 20 °C. UltraPure GdnHCl was purchased from Invitrogen. Concentrations of GdnHCl were verified using refractometry (72).

Temperature-induced unfolding transitions were generated by measuring CD or fluorescence in 2 °C increments Samples were allowed to equilibrate for two minutes at each temperature before signal measurement. The signal at each temperature was then averaged for 30 s. Reversibility was assessed by cooling samples to 25 °C after thermal denaturation and comparing CD or fluorescence emission spectra to those collected immediately prior to thermal unfolding.

GdnHCl-induced unfolding of cDHFR was monitored by CD at 222 nm, signal averaging for 30 s at each GdnHCl concentration. Unfolding of cNTL9, cSH3, and cSH2 was monitored by tryptophan fluorescence on an Aviv Model 107 ATF. Fluorescence was measured using a 280 nm excitation and either a 332 nm (cSH2 and cSH3) or 348 nm (cNTL9) emission, signal averaging for 30 s at each GdnHCl concentration. For these four proteins, unfolding was found to equilibrate rapidly, so that titrations could be generated using a Hamilton automated titrator, with a 5-minute equilibration period.

Gdn-induced unfolding of cAK and cPGK was found to equilibrate on a slower timescale than the other consensus proteins, prohibiting the use of an automated titrator. Therefore, samples at each denaturant concentration were made individually and equilibrated at room temperature for 24 hours (cAK) or 5 hours (cPGK). For each sample, the CD signal at 225 nm (cAK) or 222 nm (cPGK) was averaged for 30 seconds. Melts for both proteins were collected in buffer containing, 50 mM NaCl, 0.5 mM TCEP, and 25 mM Tris-HCl (pH 8.0).

Titrations were carried out in triplicate for each protein. Thermodynamic folding/unfolding parameters were determined by fitting a two-state linear extrapolation model to the folding/unfolding curves (73).

### Steady-state enzyme kinetics

Steady-state enzyme kinetic parameters at 20 °C were determined for cDHFR (in the direction of tetrahydrofolate formation), cAK (in the direction of ADP formation), and cPGK (in the direction of 1,3-bisphosphoglycerate formation) using absorbance spectroscopy to monitor the oxidation of NAD(P)H catalyzed either directly by the consensus enzyme (cDHFR; (74), or by the activity of an enzyme that is directly coupled to the products produced by the rate-limiting activity of the consensus enzyme (cAK and cPGK; (59, 75). The absorbance at 340 nm was monitored over time after rapid addition of consensus enzyme. Steady-state velocities were determined as the initial linear slope of the time course. Because these enzymes catalyze bi-substrate reactions, we were able to obtain Michealis-Menten kinetic parameters for each substrate by varying the concentration of one substrate at a constant, saturating concentration of the other substrate.

For cDHFR assays, a concentration of 175 nM cDHFR was used in reaction buffer containing 25 mM HEPES (pH 7.5), 150 mM NaCl. For cAK assays, a concentration of 53 nM cAK was used in reaction buffer containing 50 mM HEPES (pH 7.5), 100 mM NaCl, 20 mM MgCl_2_, 1 mM phosphoenolpyruvate, 0.1 mM NADH, 10 units pyruvate kinase, 10 units lactate dehydrogenase. For cPGK assays, a concentration of 53 nM consensus PGK was used in reaction mixture containing 100 mM Tris-HCl (pH 8.0), 3 mM MgCl_2_, 0.1 mM NADH, 5 units glyceraldehyde phosphate dehydrogenase.

For cDHFR and cPGK, k_cat_ values were measured at high temperatures using the absorbance spectroscopic assays described above at saturating concentrations of both substrates. Samples were allowed to equilibrate at the desired temperature for 5 minutes before initiation of the reaction. Consensus DHFR activity was measured up to 50 °C. Consensus PGK activity was measured up to 40 °C, the temperature of onset of denaturation of the coupling enzyme (glyceraldehyde phosphate dehydrogenase). For cAK, enzyme activity at various temperatures was measured using a direct ^31^P NMR assay previously used for an AK from *A. aeolicus* (See SI Appendix, Supplementary methods; (43). Consensus AK activity was measured up to 70 °C.

## Acknowledgements

The authors thank Ananya Majumdar for assistance in collecting and discussions of the NMR experiments. We thank the Johns Hopkins University Biomolecular NMR Center, Center for Molecular Biophysics, and Chemistry NMR Core Facility for providing facilities and resources. We thank Michael Harms for suggesting the BacDive database as a resource to classify bacterial growth temperatures. This work was supported by National Institutes of Health grant GM068462 to DB. MS was supported by NIH grants T32GM008403 and F31GM128295.

## Footnotes

† These properties include rates of folding, unfolding, degradation, compartmentalization, oligomerization, and solubility.

‡ Although for most families this search resulted in five or more free energy values, providing a good representation of the average stability, all of these stabilities may be biased by experimental constraints (expression, solubility, and baseline-resolved folding transitions.

‡ Though the 5% estimate of thermophilic sequences is likely to undercount the number of thermophilic sequences in our MSA, since not all sequences could be unambiguously assigned to a source organism, the total number of thermophiles in our MSAs is not likely to exceed 16% (the ratio of identified thermophiles to thermophiles plus mesophiles).

