## Supporting information appendix for "Consensus sequence design as a general strategy to create hyperstable, biologically active proteins"

**This PDF file includes:**

Supplementary methods

Figs. S1 to S15

Tables S1 to S5

References for SI reference citations

**Supplementary methods**

**Cloning, expression, and purification of consensus proteins**

Genes encoding the consensus sequences for each protein family were synthesized by GeneArt (ThermoFischer Scientific) as linear, double-stranded fragments. Gene fragments were designed to include a 5’ extension (5’- TAAGAAGGAGATATACATATGGGA -3’) for cloning into a modified pET24 vector, the consensus protein sequence open reading frame, a sequence encoding for a C-terminal 6x His-tag, three stop codons, and a 3’ extension (extending from the 3’ end of the stop codon sequence 5’- GGATCCAGACGTAAGCGCACC -3’) for cloning. Gene fragments were cloned into linearized vector between *NdeI* and *BamHI* restriction sites. Amino acid sequences for the resulting consensus protein constructs (including additions for cloning and purification) are shown in Table S2.

Consensus proteins were expressed in *E. coli* BL21(DE3) cells. Cells were grown in Luria broth with 50 μg/mL kanamycin at 37 °C until OD_600_ = 0.6-0.8, and were then induced with 1 mM IPTG. Cells were allowed to grow for a further 4-6 hours at 37 °C, pelleted by centrifugation, and cell pellets were stored at -80 °C.

Consensus NTL9, SH3, HD, SH2, and PGK were purified using a common protocol. Cell pellets were resuspended in either 50 mM NaPO_4_ buffer (pH 7.0; NTL9, SH3, HD, and SH2) or buffer containing 50 mM Tris-HCl (pH 8.0) and 1 mM TCEP (PGK) with a Pierce EDTA-free protease inhibitor cocktail tablet (Thermo Scientific). Cells were lysed by sonication. Cell lysate was centrifuged to separate soluble and insoluble fractions, from which the supernatant was collected. Proteins were purified using Ni-NTA chromatography followed by ion-exchange chromatography. Purified proteins were dialyzed in either 25 mM NaPO_4_ (pH 7.0) and 150 mM NaCl (NTL9, SH3, HD, and SH2) or 25 mM Tris-HCl (pH 8.0), 50 mM NaCl, and 0.5 mM TCEP (PGK).

Consensus DHFR and AK were purified denatured in urea, since these proteins prepared under native conditions were found to retain endogenous substrates from the cells. Cell pellets were resuspended in buffer containing either 50 mM NaPO_4_ (pH 7.0) and 8 M urea (DHFR) or 50 mM Tris-HCl (pH 8.0), 1 mM TCEP, and 8 M urea (AK). Cells were lysed by sonication, centrifuged to separate soluble and insoluble portions. The soluble fractions containing denatured AK and DHFR were loaded onto Ni-NTA columns and were refolded on the column by washing into buffer without urea. Proteins were eluted under native conditions and further purified by ion exchange chromatography. Purified consensus DHFR was dialyzed 25 mM NaPO_4_ (pH 7.0) and 150 mM NaCl. Purified consensus AK was dialyzed into 25 mM Tris-HCl (pH 8.0), 50 mM NaCl, and 0.5 mM TCEP. All proteins were frozen in liquid nitrogen and stored at -80 °C. Protein concentrations for all experiments were determined by UV-Vis absorbance spectroscopy (1).

**NMR spectroscopy**

^15^N- and ^13^C,^15^N-isotopically labeled proteins were expressed and purified in *E. coli* BL21(DE3) cells grown in M9 minimal media supplemented with ^15^NH_4_Cl and either ^12^C- or ^13^C-glucose (Cambridge Isotope Laboratories). Proteins were purified as described above, and were concentrated to 400-800 μM.

^1^H-^15^N HSQC spectra were collected for ^15^N-labeled consensus NTL9, SH3, HD, SH2, DHFR, and AK at 25 °C on Bruker Avance or Avance II 600 MHz spectrometers equipped with cryoprobes. NMR samples for NTL9, SH3, HD, and SH2 contained 150 mM NaCl, 5% D_2_O, and 25 mM NaPO_4_ (pH 7.0). Spectra of consensus DHFR were collected under the same conditions both without and with a 1:1 molar equivalent of methotrexate (Sigma Aldrich). Spectra of consensus AK were collected in 50 mM NaCl, 1 mM TCEP, 5% D_2_O, and 25 mM Tris-HCl (pH 7.5) at 25 °C. For ^15^N-labeled consensus PGK, ^1^H-^15^N TROSY spectra were collected on a Varian 800 MHz spectrometer at 35 °C. NMR samples of consensus PGK contained 50 mM NaCl, 1 mM TCEP, 5% D_2_O, and 25 mM Tris-HCl (pH 7.5).

All data were processed and analyzed using NMRPipe (2) and NMRFAM-SPARKY (3). Backbone assignments for consensus NTL9, SH3, and SH2 were made using standard triple-resonance experiments including HNCACB, HNCO, HN(CA)N on Bruker Avance or Avance II 600 MHz spectrometers. Backbone chemical shift data were used for secondary structure predictions using TALOS-N (4).

Consensus SH3 peptide binding was monitored using ^1^H-^15^N HSQC spectra as described in the main text. At each peptide concentration, chemical shift perturbations for each residue (Δδ_NH,i_) were calculated as a weighted Pythagorean distance:

$\text{∆δ}_{\text{NH,i}}\text{=}\sqrt{\frac{\text{1}}{\text{2}}{\text{[}\text{∆}\text{δ}}_{\text{H,i}}^{\text{2}}\text{ + 0.14}{\text{∆}\text{δ}}_{\text{N,i}}^{\text{2}}\text{]}}$	(1)

where Δδ_X,i_ is the change in chemical shift of the *i*^th^ resonance in either ^15^N or ^1^H, relative to the apo-protein chemical shift value (5). The weighting factor of 0.14 accounts for differences in ^1^H and ^15^N chemical shift sensitivities. For the ten peaks showing the greatest changes, chemical shift perturbations at each peptide concentrations were globally fit to the single-site binding equation:

${\text{ }\text{∆}\text{δ}}_{\text{NH}}\text{ }\text{=}{\text{ }\text{∆}\text{δ}}_{\text{max}}\frac{\left( \left[ \text{P} \right]_{\text{t}}\text{ }\text{+}\left[ \text{L} \right]_{\text{t}}\text{ }\text{+}{\text{ }\text{K}}_{\text{d}} \right) -\text{ }\sqrt{\left( \left[ \text{P} \right]_{\text{t}}\text{ }\text{+}\text{ }\left[ \text{L} \right]_{\text{t}\text{ }}\text{+}\text{ }\text{K}_{\text{d}} \right)^{\text{2}}-\text{ }\text{4}\left[ \text{P} \right]_{\text{t}}\left[ \text{L} \right]_{\text{t}}}}{\text{2}\left[ \text{P} \right]_{\text{t}}}$	(2)

where Δδ_max_ is a local parameter for maximal chemical shift perturbation for each peak, [P]_t_ is the total cSH3 concentration, [L]_t_ is the total peptide concentration, and K_d_ is a global dissociation constant (5). In the fit, Δδ_max_ values were optimized locally (for each of the ten resonances), K_d_ was optimized globally, and [P]_t_ was fixed at its known value.

Consensus AK enzyme activity at various temperatures was measured using a direct ^31^P NMR assay previously used for an AK from *A. aeolicus* (6). Conversion of ADP to ATP and AMP was monitored in real time by ^31^P NMR using a Bruker Avance III HD 400 MHz spectrometer equipped with a broadband probe. ADP at an initial concentration of 13 mM was rapidly mixed with cAK, and 1D ^31^P NMR spectra were collected continuously (32 scans per spectrum with an inter-scan delay of 8-10 seconds) until equilibrium was reached. For ATP and ADP , peak areas of each ^31^P resonance (three for ATP, two for ADP) were averaged and were fitted along with the single AMP peak to obtain forward and reverse rate constants. Forward rate constants were converted to k_cat_ values by dividing by cAK concentration. To maintain the kinetics in a measurable range, we decreased the enzyme concentration as temperature was increased from 1.1 μM at 20 °C to 24 nM at 70 °C.

**Sequence analysis**

Analysis of curated multiple sequence alignments and consensus sequences was performed using in-house scripts (available upon request). Residues were sorted into groups based on physiochemical properties: charged residues (K, R, D, E), polar uncharged residues (N, C, Q, S, T, H), and nonpolar residues (A, I, L, M, V, F, W, G, P, Y). Sequence net charge was calculated assuming contributions of +1 for all K and R residues, -1 for all E and D residues, and 0 for H residues.

Sequence entropies for positions in the multiple sequence alignment were calculated as described in the main text. In all position-by-position comparisons to naturally-occurring proteins, only positions represented in the consensus sequence were considered.

For a particular sequence feature *f*, the position the value of the feature for the consensus sequences (*f_c_*) within distributions for multiple sequence alignments (*f_MSA_*) were evaluated using a Z-score,

(3)

where *s_MSA_* is the standard deviation of *f_MSA_* values. In this context, the Z-score can be thought of as the number of standard deviations the consensus sequence is from the average from the multiple sequence alignment (*<f_MSA_>*).

Homology models for all consensus proteins were made using SWISS-MODEL(7), using the structure of the sequence displaying the highest sequence identity to the consensus sequence as a template. Residue-specific solvent accessible surface areas were calculated from these homology models using GETAREA (8). Surface, intermediate, and buried positions were defined as positions at which residues show greater than a 50%, 20%--50%, and less than 20% changes in side-chain solvent accessible surface area in homology models of the consensus proteins relative to an ensemble of Gly-X-Gly tripeptides as calculated by the GETAREA algorithm.

To determine the number of thermophilic/mesophilic sequences in the MSAs composed of predominantly bacterial sequences (NTL9, DHFR, AK, and PGK), the source organism of each sequence was identified from the database-specific sequence IDs using the UniProt database (9). For the subset of sequences for which a source organism could be identified by UniProt (on average 97% of sequences in the MSAs) and that belonged to the bacteria or archaea domains, we used the BacDive database (accessed on 3/25/19) to determine whether the source bacterial or archael organism was a thermophile or mesophile using the "Temperature range" classification in the “Culture and growth conditions” data fields under the “Advanced search” (10). Only organisms that contained both genus and species information in BacDive (10381 sequences total) were assigned to the thermophilic/hyperthermophilic or mesophilic classifications to limit ambiguous classifications. Of these sequences, 853 were thermophilic or hyperthermophilic and 9528 were mesophilic organisms were identified by the BacDive database.

A

B

C

D

E

F

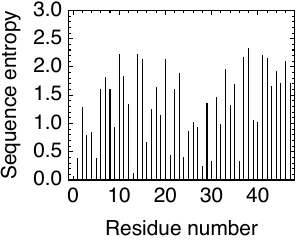

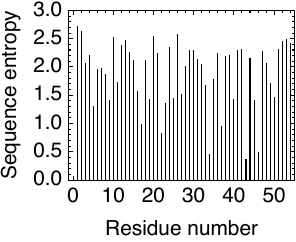

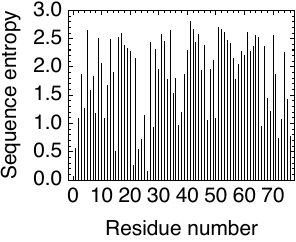

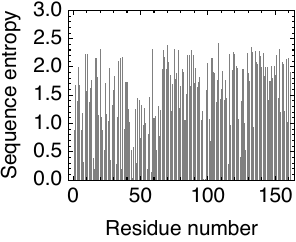

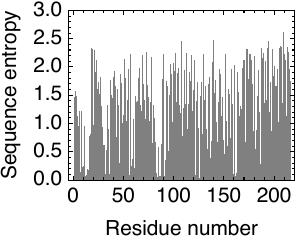

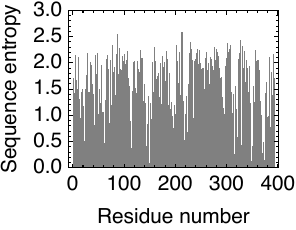

Figure S1. Position-specific conservations of multiple sequence alignments. Sequence entropies shown for all consensus positions in multiple sequence alignments of (A) NTL9, (B) SH3, (C) SH2, (D) DHFR, (E) AK, and (F) PGK.

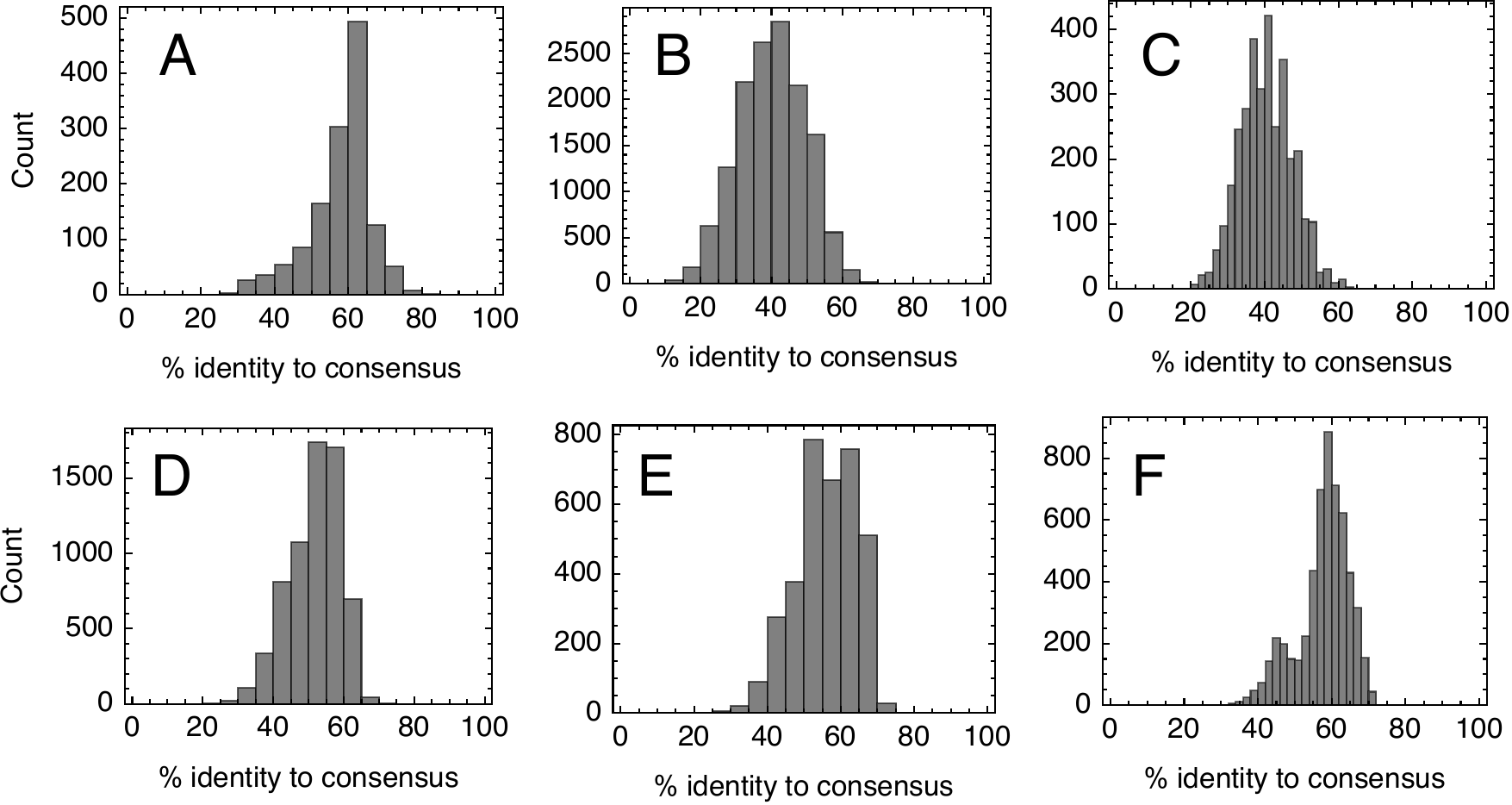

Figure S2. Identities of naturally-occurring sequences to consensus sequence for (A) NTL9, (B) SH3, (C) HD, (D) SH2, (E) DHFR, (F) AK, and (G) PGK.

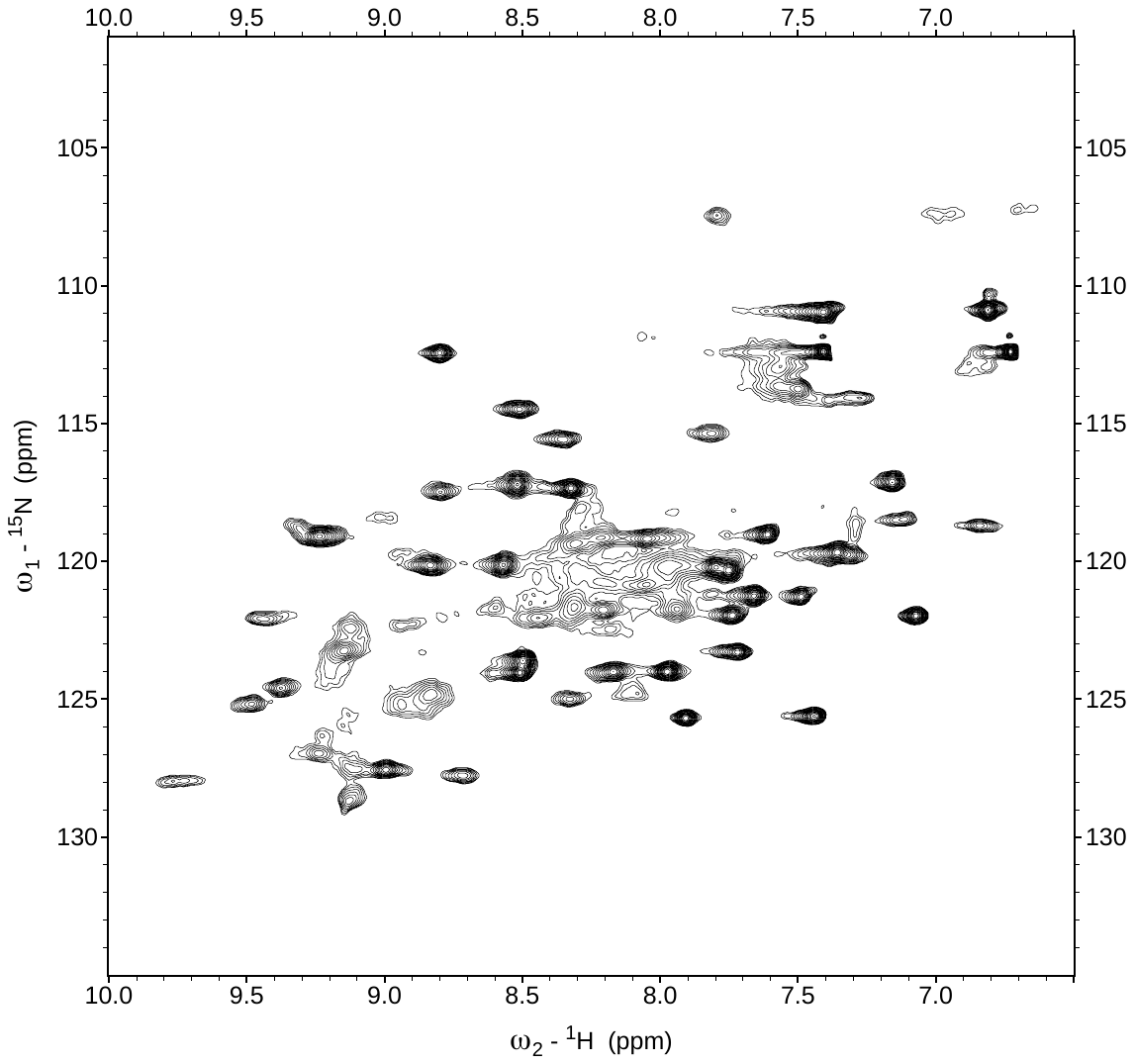

Figure S3. ^1^H-^15^N HSQC spectrum of cDHFR in the apo state at 600 MHz. Experimental conditions: 25 mM sodium phosphate, 150 mM sodium chloride, pH 7.0, 25 °C.

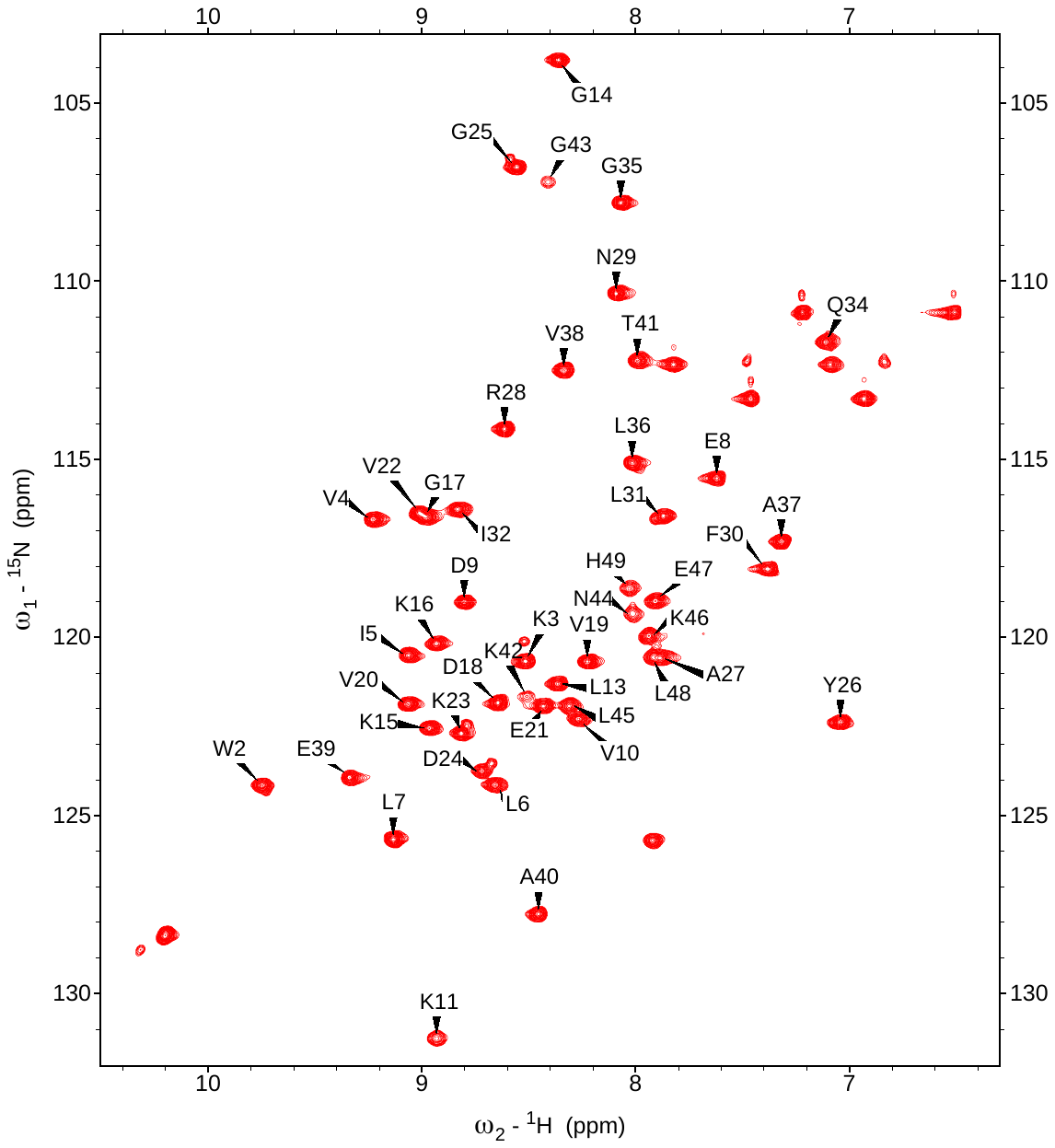

Figure S4. ^1^H-^15^N HSQC spectrum of cNTL9 at 600 MHz. Assigned peaks are label. Experimental conditions: 25 mM sodium phosphate, 150 mM sodium chloride, pH 7.0, 25 °C.

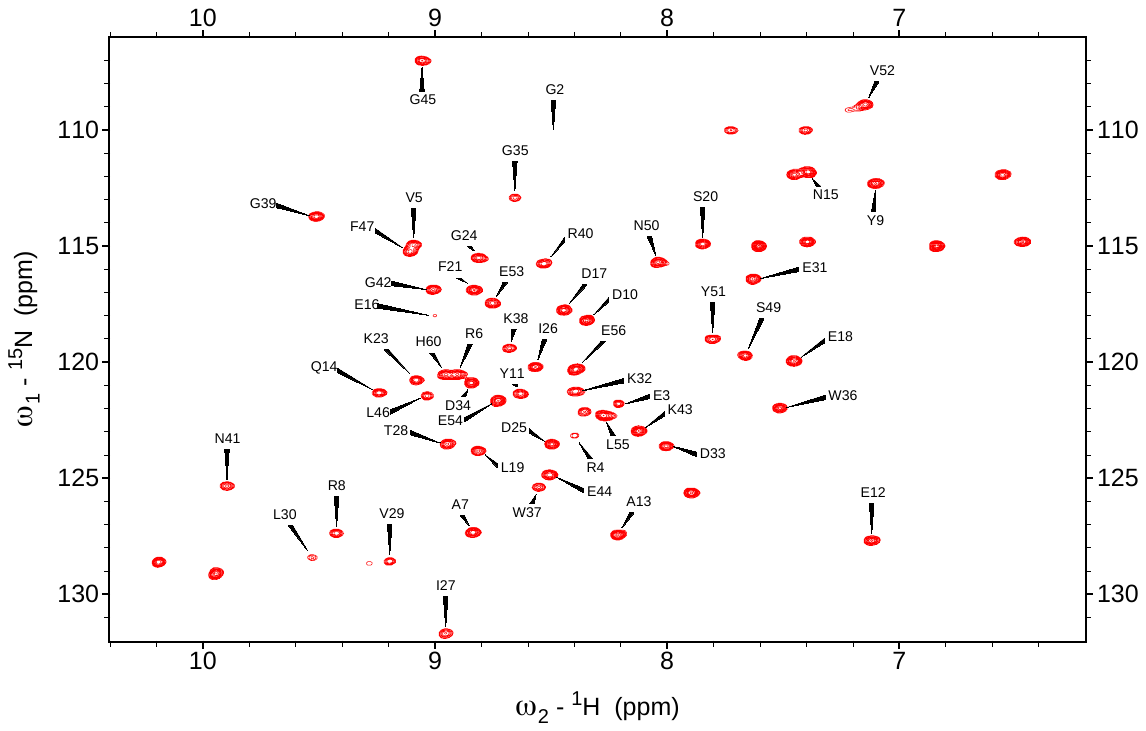

Figure S5. ^1^H-^15^N HSQC spectrum of cSH3 at 600 MHz. Assigned peaks are label. Experimental conditions: 25 mM sodium phosphate, 150 mM sodium chloride, pH 7.0, 25 °C.

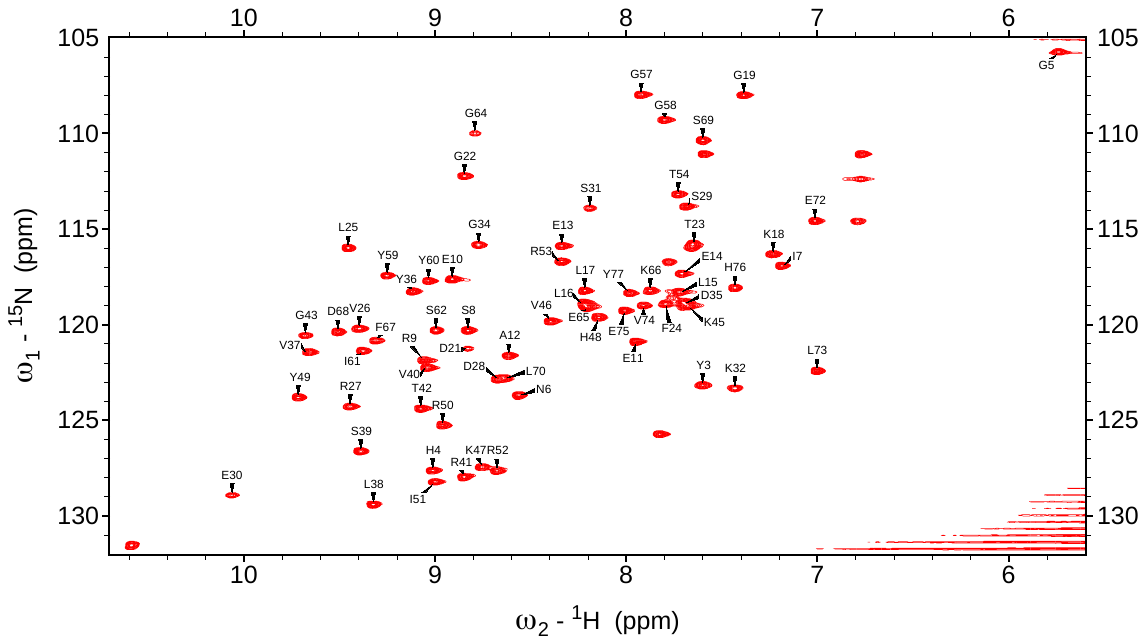

Figure S6. ^1^H-^15^N HSQC spectrum of cSH2 at 600 MHz. Assigned peaks are label. Experimental conditions: 25 mM sodium phosphate, 150 mM sodium chloride, pH 7.0, 25 °C.

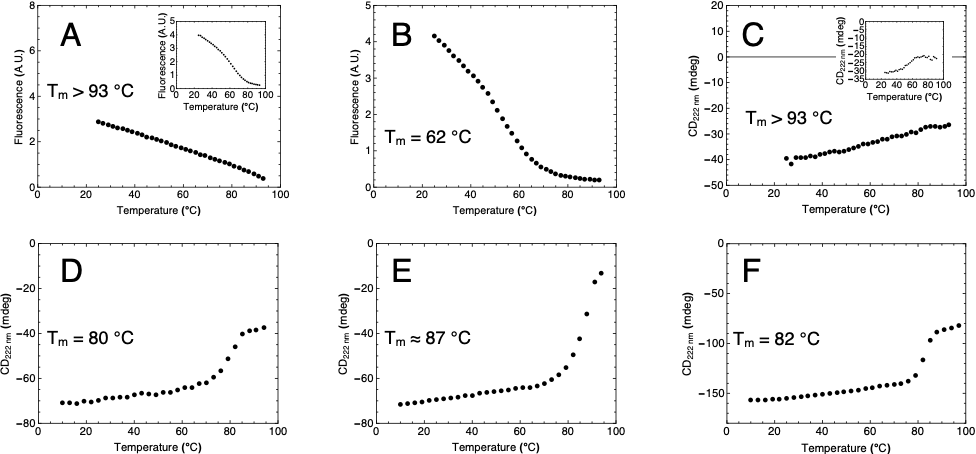

**Figure S7.** Temperature-induced unfolding curves for (A) cNTL9, (B) cSH3, (C) cSH2, (D) cDHFR, (E) cAK, and (F) cPGK. For cNTL9 and cSH2 (A and C), there are no obvious thermal unfolding transitions up to 93 °C; insets show unfolding transitions in the presence of 4 and 2 M GdnHCl, respectively. Plots show raw values measured using circular dichroism (SH2, DHFR, AK, and PGK) or fluorescence (NTL9, and SH3) spectroscopies.

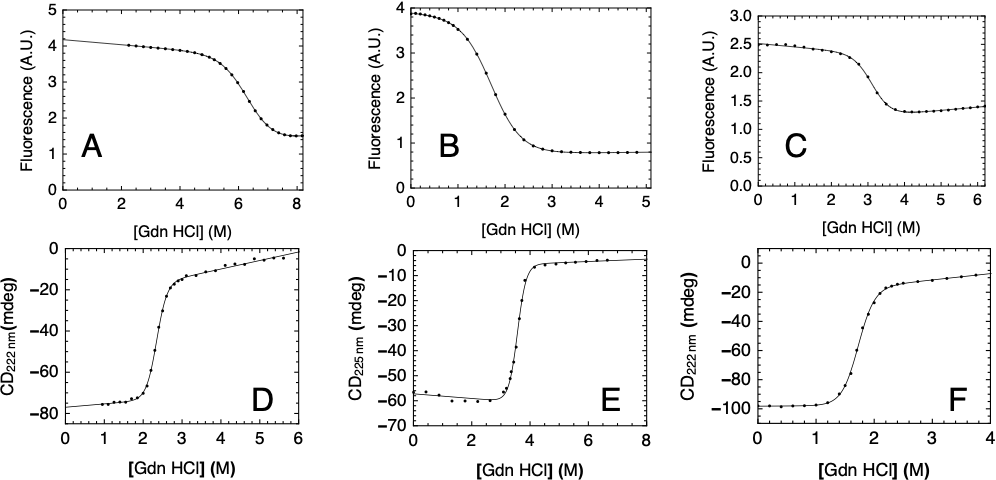

**Figure S8.** Guanidine hydrochloride-induced unfolding curves for (A) cNTL9, (B) cSH3, (C) cSH2, (D) cDHFR, (E) cAK, and (F) cPGK. Plots show raw values measured using circular dichroism (cDHFR, cAK, and cPGK) or fluorescence (cNTL9, cSH3, cSH2) spectroscopies. Solid lines are obtained from fitting a two-state model to the data. Parameters obtained from two-state fits to raw data were used to convert to fraction folded. Experimental conditions are as noted in main text.

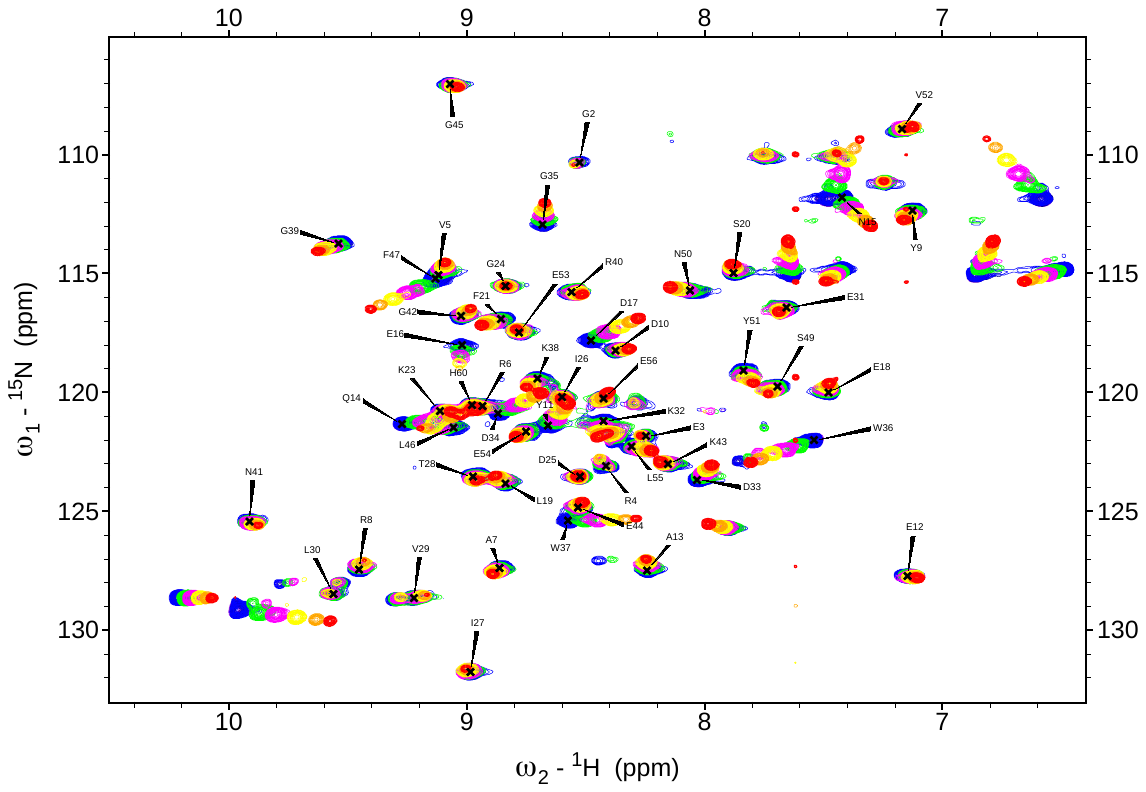

**Figure S9.** ^1^H-^15^N HSQC peptide titration of cSH3. Experiments were collected at 0- (purple), 0.05- (dark blue), 0.125- (blue), 0.5- (dark green), 1.25- (green), 2.5- (magenta), 5- (yellow), 10- (orange), and 20-fold (red) saturation of nonisotopically labeled peptide concentrations. 200 μM consensus SH3 was used for all experiments. Residues assigned in apo-spectrum are labeled.

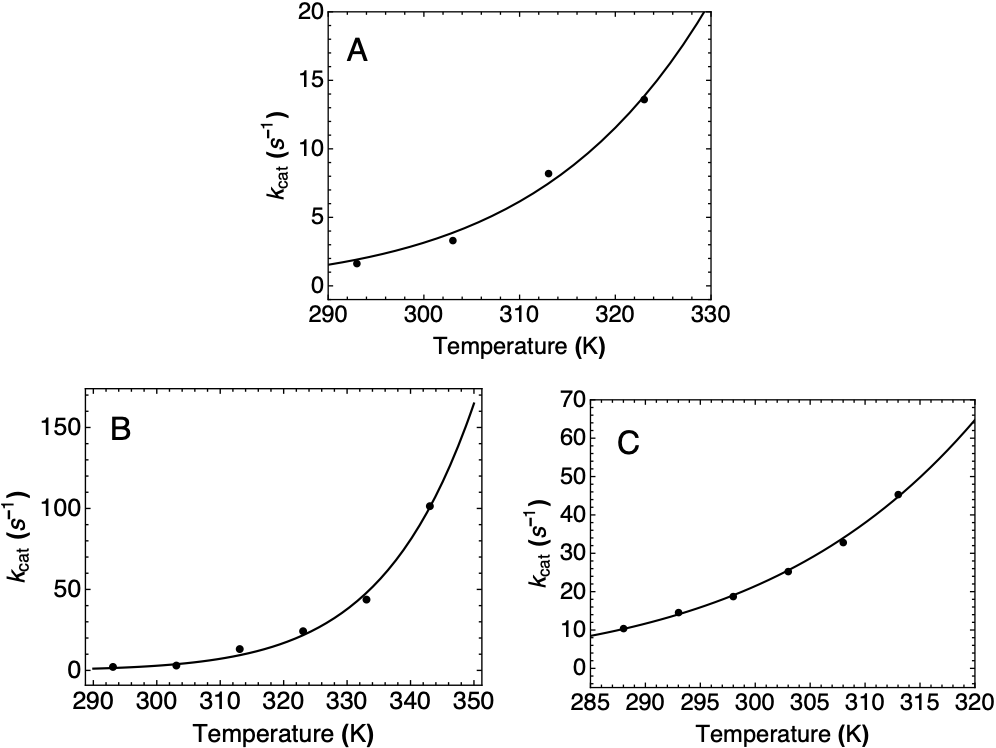

**Figure S10.** Temperature dependence of (A) cDHFR (in the DHF + NADPH to THF + NADP^+^ direction), (B) cAK (in the 2ADP to ATP + AMP direction), and (C) cPGK (in the 3-PG + ATP to 1,3-BPG + ADP direction). Activities for cDHFR andc PGK were measured using absorbance spectroscopic assays; activities for cAK were measured using ^31^P NMR assay (see Supplementary methods, main text). Solid lines are obtained by fitting an Arrhenius model to the data. Experimental conditions are given in the main text.

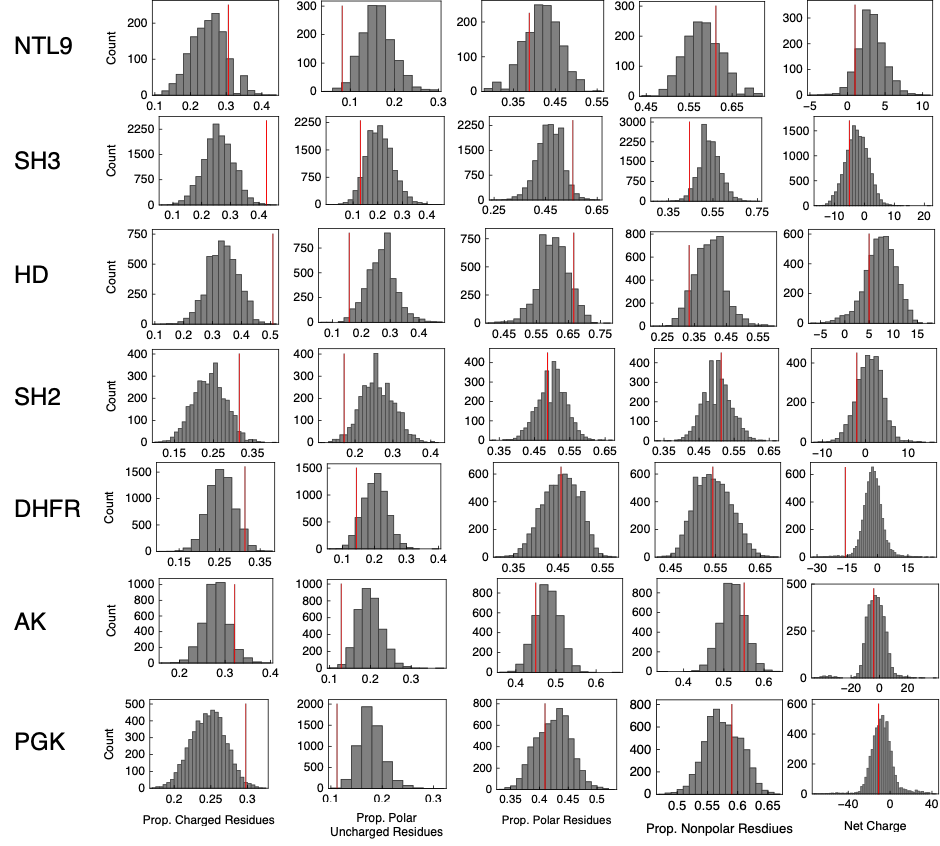

**Figure S11**. Sequence biases of consensus sequences. Distributions of proportions of sequence made up of charged, polar uncharged, total polar, and nonpolar residues, and net charge of extant sequences in final multiple sequence alignments. Red lines indicate where the consensus sequence lies for each parameter.

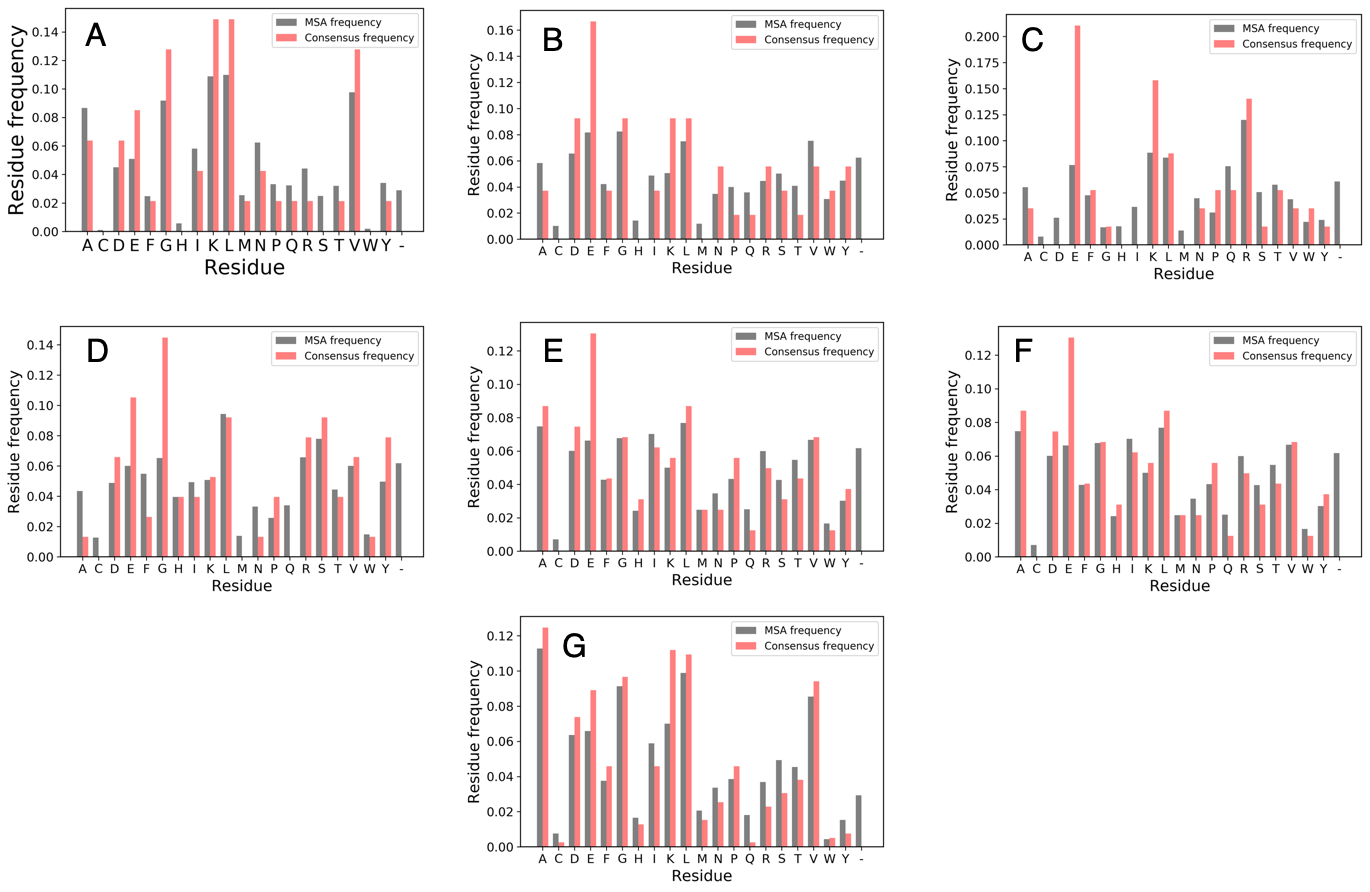

**Figure S12.** Residue frequencies for extant sequences in MSA (black) and consensus sequence (red) for (A) NTL9, (B) SH3, (C) HD, (D) SH2, (E) DHFR, (F) AK, and (G) PGK.

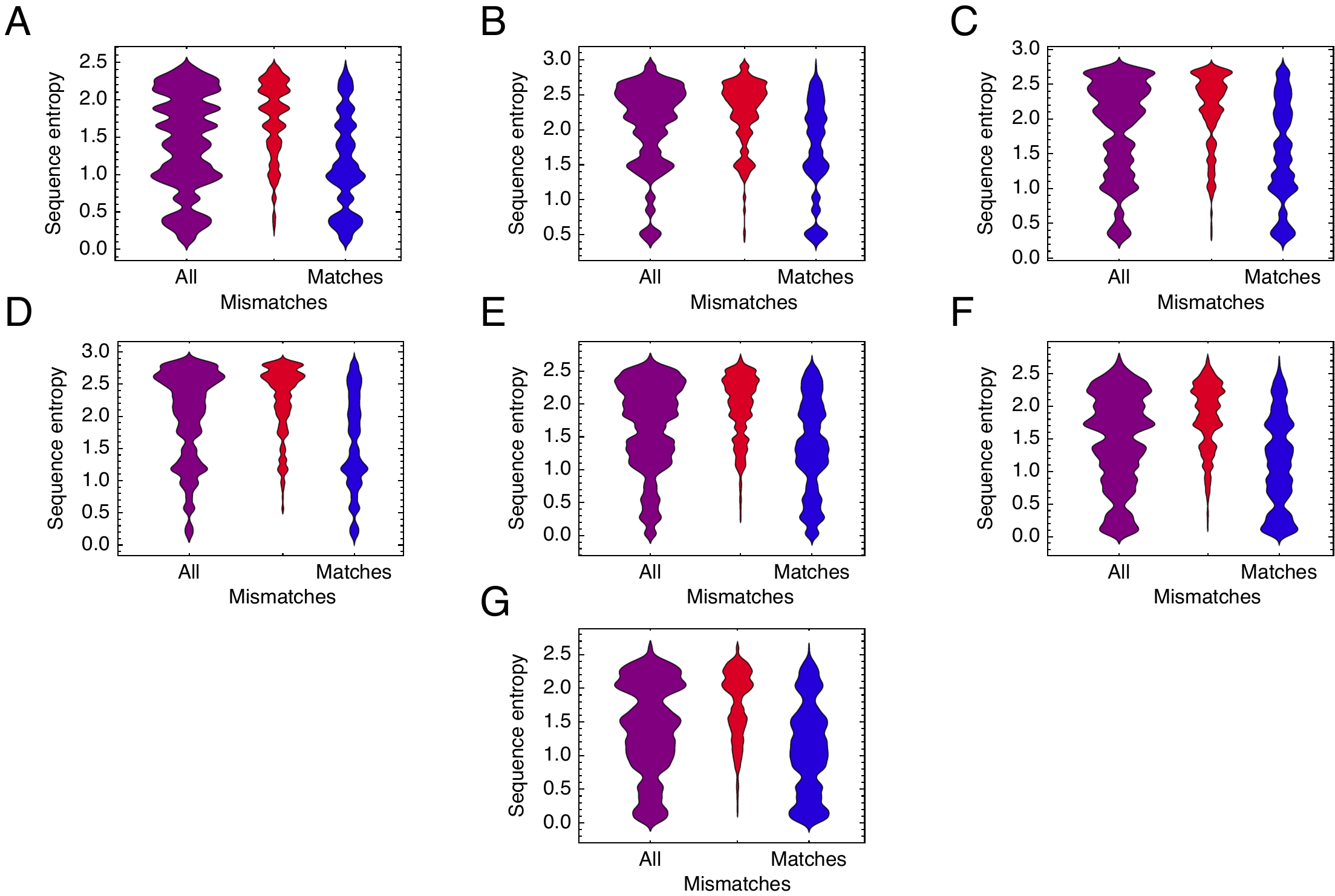

**Figure S13**. Positional conservation bias of consensus mismatches. Distributions of sequence entropy values for all positions in MSA (“all”; purple), positions at which extant sequences differ from the consensus sequence (“mismatches”; red), and positions at which extant sequences match the consensus sequence (“matches”; blue) for consensus (A) NTL9, (B) SH3, (C) HD, (D) SH2, (E) DHFR, (F) AK, and (G) PGK.

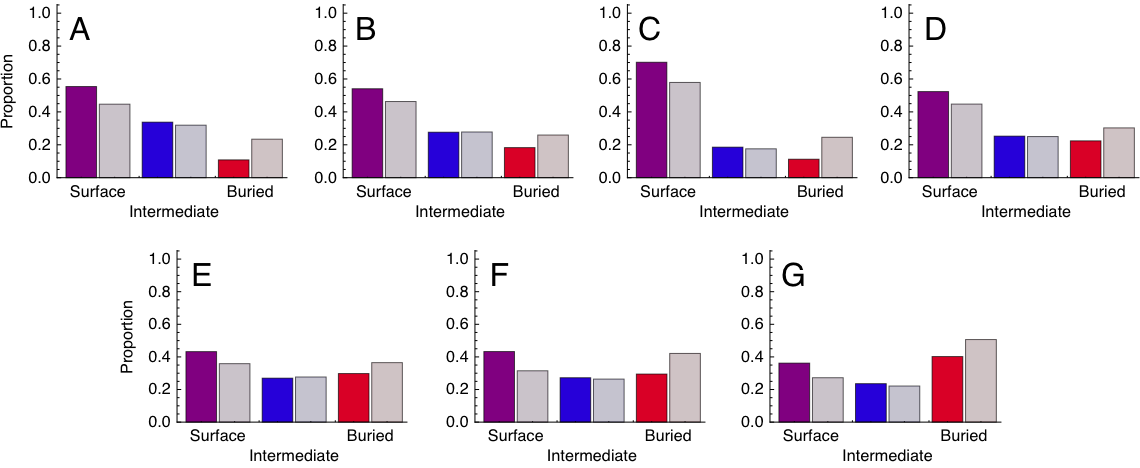

**Figure S14**. Surface exposure of consensus mismatches. Dark bars show the proportion of mismatch residues that are at surface (purple), intermediate (blue), and buried positions (red) for different consensus protein targets. Light bars show the proportion of all residues at surface, intermediate, and buried positions. (A) NTL9, (B) SH3, (C) HD, (D) SH2, (E) DHFR, (F) AK, and (G) PGK. The degree of burial is determined as described in the main text.

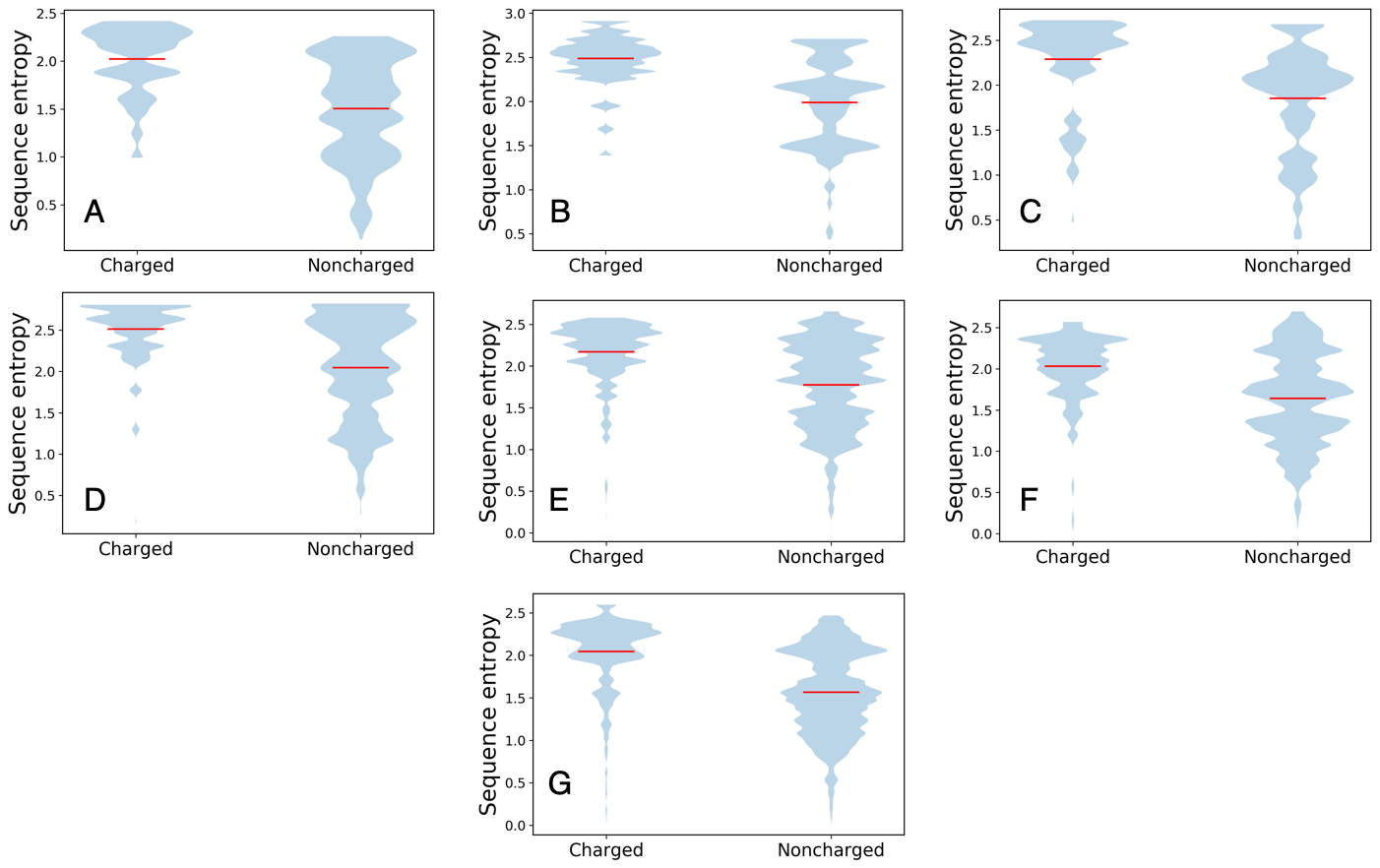

**Figure S15.** Sequence entropy distributions of consensus substitutions to from uncharged to charged residues (left) and substitutions among uncharged residues (right) for (A) NTL9, (B) SH3, (C) HD, (D) SH2, (E) DHFR, (F) AK, and (G) PGK. The red line signifies the mean of each distribution.

| Table S1. Data for sequence sets used for consensus sequence generation. | | | | | |
| --- | --- | --- | --- | --- | --- |
| Protein family | Database  Accession identity  Date accessed | Number of sequence in initial set | Number of sequences after curation | Average pairwise sequence identity | Phylogenetic distribution of sequences |
| NTL9 | Pfam  PF01281  5/20/16 | 1911 | 1355 | 42% | Ar: 0%  Bac: 83.9%  Eu: 16.1% |
| SH3 | SMART  SM00326  4/4/17 | 54382 | 14474 | 26% | Ar: 0.0%  Bac: 4.3%  Eu: 95.7% |
| SH2 | Pfam  PF00017  5/20/16 | 10051 | 3326 | 28% | Ar: 0.0%  Bac: 0.4%  Eu: 99.6% |
| DHFR | InterPro  IPR001796  5/2/17 | 16038 | 6542 | 37% | Ar: 2.3%  Bac: 92.4%  Eu: 5.3% |
| AK | InterPro  IPR007862  2/9/17 | 13905 | 3534 | 43% | Ar: 4.1 %  Bac: 74.5%  Eu: 21.5% |
| PGK | InterPro  IPR001576  4/5/17 | 17724 | 5581 | 45% | Ar: 6.2%  Bac: 82.7%  Eu: 11.1% |
| Databases used to obtain sequence sets for each protein family are noted along with the database-specific accession identity of each protein family and the date sequence sets were obtained. The number of sequences in the initial set gathered directly from the database is reported along with the number of sequences in the final set used for consensus sequence generation after sequences were removed based on sequence length and sequence identity (see main text). Reported average pairwise identity is the average of the identities of all pairwise comparisons between sequences in the final sequence set used. Phylogenetic distributions represent the percentage of sequences in the final sequence set classified as archeal (Ar), bacterial (Bac), and eukaryotic (Eu) by the database used to obtain the sequence set. | | | | | |

| Table S2. Consensus sequence constructs generated for protein families. | |
| --- | --- |
| Protein family | Consensus sequence |
| NTL9 | MGWKVILLEDVKGLGKKGDVVEVKDGYARNFLIPQGLAVEATKGNLKELHHHHHH |
| SH3 | MGERVRARYDYEAQNEDELSFKKGDIITVLEKDDGWWKGRNGKEGLFPSNYVEELEHHHHHH |
| SH2 | MGWYHGNISREEAEELLLKGPDGTFLVRDSESKPGDYVLSVRTGGKVKHYRIRRTDGGGYYISGGEKFDSLPELVEHYHHHHHH |
| DHFR | MGISLIVAVAENGVIGKDNDLPWHLPEDLKHFKELTMGHPVIMGRKTFESIGRPLPGRRNIVLTRDPDYQAEGAEVVHSLEEALALAKEAEEVFVIGGAEIYAQALPLADRLYLTEIDADFEGDTFFPEIDSEWKEVSREEHPADEKNGYDYTFVTYERKKHHHHHH |
| AK | MGWRIILLGPPGAGKGTQAKRIVEKYGIPHISTGDMLRAAIKAGTELGKKAKSYMDAGELVPDEIVIGLVKERLAQPDCNGFLLDGFPRTIPQAEALDELLKELGVKLDAVIELDVPDEELVERLSGRRVCPAKCGRTYHVKFNPPKVEGVCDVCGEELIQRDDDKEETVRKRLEVYHEQTAPLIDYYKKKGLLVTVDGTGSIDEVFADILAALGKKKHHHHHH |
| PGK | MGNKTIDDLDLKGKRVLVRVDFNVPLKDGKITDDTRIRAALPTIKYLLEKGAKVILMSHLGRPKGEVDPKFSLAPVAKRLSELLGKPVKFADDCVGEEAEAAVAALKPGEVLLLENLRFHKGEEKNDPEFAKKLASLGDVYVNDAFGTAHRAHASTVGVAKFLPAAAGFLMEKELEALGKALENPERPFVAILGGAKVSDKIGVIENLLDKVDKLIIGGGMANTFLKAQGYEVGKSLVEEDKLDTAKELLEKAKEKGVKIVLPVDVVVADEFSADAETKVVPVDEIPDDWMGLDIGPKTVELFAEAIKDAKTIVWNGPMGVFEFEPFAKGTKAVAKAIAEATGAFSIVGGGDTAAAVNKLGLADKFSHISTGGGASLEFLEGKELPGVAALEDKHHHHHH |
| Consensus sequences generated for each protein family are shown. Consensus sequence derived from multiple sequence alignments are shown in black. Residues added to consensus sequences for cloning (N-terminal MG), quantification (N-terminal W, added if consensus sequence did not contain a tryptophan), or purification (C-terminal His-tag) purposes are shown in red. | |

| **Table S3. Stabilities of naturally-occurring proteins reported in literature.** | | | | | | |
| --- | --- | --- | --- | --- | --- | --- |
| Protein | Organism | Denaturant used | ΔG°_H2O_  (kcal/mol) | Mean  ΔG°_H2O_  (kcal/mol) | *m*-value  (kcal/mol/M) | Reference |
| **NTL9** | Consensus | Gdn HCl | -7.9 | -3.2 | 1.23 | This study |
|  | *E. coli* | Gdn HCl | -1.98 |  | 1.21 | (11) |
|  | *B. stearothermophilus* | Urea | -4.5 |  | NR | (12) |
| **SH3** | Consensus | Gdn HCl | -3.2 | -3.5 | 1.88 | This study |
|  | Human Fyn | Gdn HCl | -5.0 |  | 1.5 | (13) |
|  | Human BTK | Gdn HCl | -2.6 |  | 1.18 | (14) |
|  | *C. elegans* Sem-5 | Gdn HCl | -4.1 |  | 1.7 | (15) |
|  | Human Src | Gdn HCl | -4.1 |  | 1.6 | (16) |
|  | *D. melanogaster* drk | Gdn HCl | -2.18 |  | 1.39 | (17) |
|  | Human PI3K | Gdn HCl | -3.23 |  | 2.33 | (18) |
|  | Chicken α-spectrin | Urea | -3.6 |  | 0.74 | (19) |
|  | Yeast Abp1p | Gdn HCl | -3.1 |  | 1.64 | (20) |
|  | Mouse Crk-II | Urea | -3.33 |  | 0.66 | (21) |
| **HD** | Consensus | Gdn HCl | -8.1 | -2.8 | 1.6 | (22) |
|  | *D. melanogaster* Engrailed | Gdn HCl | -3.62 |  | 1.39 | (22) |
|  | Rat TTF-1 | Urea | -1.62 |  | NR | (23) |
|  | Human Hesk-1 | Urea | -4.43 |  | 0.9 | (24) |
|  | Rat ISL-1 | Urea | -1.67 |  | 1.17 | (25) |
|  | *D. melanogaster* Antp | Gdn HCl | -2.85 |  | 0.61 | (26) |
|  | *D. melanogaster* TTF-1 | Gdn HCl | -2.59 |  | 0.65 | (26) |
| **SH2** | Consensus | Gdn HCl | -7.2 | -3.6 | 2.28 | This study |
|  | Human CSK | Gdn HCl | -6.57 |  | 2.19 | (27) |
|  | Human BTK | Gdn HCl | -2.95 |  | 1.34 | (28) |
|  | Human Stat5b | Gdn HCl | -1.2 |  | NR | (29) |
| **DHFR** | Consensus | Gdn HCl | -10.1 | -3.4 | 4.32 | This study |
|  | *E. coli* | Gdn HCl | -6.97 |  | 4.1 | (30) |
|  | Human | Gdn HCl | -4.2 |  | 5.87 | (31) |
|  | *M. profunda* | Urea | -1.89 |  | 1.03 | (32) |
|  | *S. benthica* DB21MT-2 | Urea | -2.08 |  | 1.1 | (33) |
|  | *S. benthica* DB6705 | Urea | -1.89 |  | 1.12 | (33) |
|  | *S. frigidmarina* | Urea | -1.98 |  | 0.96 | (33) |
|  | *S. oneidensis* | Urea | -1.6 |  | 1.4 | (33) |
|  | *S. putrefaciens* | Urea | -1.98 |  | 1.65 | (33) |
|  | *S. violacea* | Urea | -1.91 |  | 0.86 | (33) |
|  | *B. stearothermophilus* | Gdn HCl | -7.4 |  | NR | (34) |
|  | *L. casei* | Urea | -4.6 |  | 2.4 | (35) |
|  | Mouse | Urea | -4.4 |  | NR | (36) |
| **AK** | Consensus | Gdn HCl | -13.6 | -5.3 | 3.83 | This study |
|  | Yeast | Gdn HCl | -5.47 |  | 7.29 | (37) |
|  | *B. subtilis* | Gdn HCl | -3.4 |  | 2.9 | (38) |
|  | Chicken | Gdn HCl | -3.8 |  | 4.7 | (39) |
|  | Pig | Gdn HCl | -3.9 |  | 4.8 | (39) |
|  | *E. coli* | Gdn HCl | -9.8 |  | 2.9 | (40) |
| **PGK** | Consensus | Gdn HCl | -7.5 | -6.1 | 4.34 | This study |
|  | Yeast | Gdn HCl | -3.63 |  | 5.86 | (41) |
|  | *T. thermophiles* | Gdn HCl | -6.32 |  | 2.7 | (41) |
|  | *E. coli* | Urea | -9.3 |  | NR | (42) |
|  | Human PGK1 | Urea | -8.3 |  | 3.4 | (43) |
|  | Horse | Gdn HCl | -2.87 |  | NR | (44) |
| Data were gathered by searching the PubMed database for free energies of folding for naturally-occurring sequences of respective protein families. The search was limited to free energy values determined by chemical denaturation experiments and to proteins that appeared monomeric. “Mean “ folding free energy denotes average of values for extant homologues. NR denotes values that were not reported in the studies. | | | | | | |

**Table S4. Temperature dependence of consensus enzyme activities.**

|  | Temperature (°C) | k_cat_  (s^-1^) | E_act_  (kcal mol^-1^) | A_0_  (s^-1^) | Reference |
| --- | --- | --- | --- | --- | --- |
| **DHFR** |  |  |  |  |  |
| Consensus | --- | --- | 12.4 | 3.3 × 10^9^ | This study |
|  | 20 | 1.7 |  |  |  |
|  | 30 | 3.3 |  |  |  |
|  | 40 | 8.2 |  |  |  |
|  | 50 | 13.6 |  |  |  |
|  | ^a^*60* | ^a^*24.0* |  |  |  |
| *B. stearothermophilus* | 60 | 4.8 | NR | NR | (45) |
| *T. maritima* | 60 | 0.76 | ^b^11.1 | NR | (46) |
| **AK** |  |  |  |  |  |
| Consensus | --- | --- | 16.9 | 5.8 × 10^12^ | This study |
|  | 20 | 1.7 |  |  |  |
|  | 30 | 3.3 |  |  |  |
|  | 40 | 12.9 |  |  |  |
|  | 50 | 23.8 |  |  |  |
|  | 60 | 43.6 |  |  |  |
|  | 70 | 101.6 |  |  |  |
| *A. aeolicus* | 50 | ^c^500 | NR | NR | (6) |
| **PGK** |  |  |  |  |  |
| Consensus | --- | --- | 10.5 | 1.0 × 10^9^ | This study |
|  | 15 | 10.6 |  |  |  |
|  | 20 | 14.7 |  |  |  |
|  | 25 | 18.9 |  |  |  |
|  | 30 | 25.3 |  |  |  |
|  | 35 | 32.9 |  |  |  |
|  | 40 | 45.4 |  |  |  |
|  | ^a^*60* | ^a^*128* |  |  |  |
|  | ^a^*70* | ^a^*203* |  |  |  |
|  | ^a^*75* | ^a^*254* |  |  |  |
| *T. maritima* | 40 | 231 | 9.1 | NR | (47) |
| *T. brockii* | 40 | *^c^*600 | NR | NR | (48) |
| *M. fervidus* | 60 | 150 | 9.80 | NR | (49) |
| *P. woesei* | 70 | 113 | 16.7 | NR | (49) |
| *T. thermophilus* | 75 | 1015 | NR | NR | (50) |

Temperature dependence of consensus enzyme activities. DHFR (in direction of tetrahydrofolate formation) and PGK (in the direction of 1,3-bisphosphoglycerate formation) turnover numbers (k_cat_) were measured using absorbance spectroscopy. AK (in the direction of ATP and AMP formation) turnover numbers were measured using a real-time ^31^P NMR . Activation energies (E_act_) and pre-exponential factors (A_0_) were determined by fitting an Arrhenius model to the data (Figure S10). NR denotes that values were not reported in the studies.

*^a^*denotes k_cat_ values that were extrapolated using fitted parameters in an Arrhenius model

*^b^*Arrhenius plot shows two linear regimes with different slopes with a break point around 25 °C . The E_act_ value reported is from the linear regime above 25 °C.

*^c^*denotes values that were estimated from graphs in references

| Table S5. Thermophilic/mesophilic sequence composition for predominantly bacterial protein families. | | | | |
| --- | --- | --- | --- | --- |
| Protein family | Bacterial + archeal sequences in set | Sequence source organisms identified by BacDive | Thermophilic (T),  mesophilic (M), and  unclassified (U) sequences | Average identity to consensus for thermophilic (T) and mesophilic (M) sequences |
| NTL9 | 1136 | 761 | T: 66 (4.8%)  M: 510 (37.6%)  U: 185 | T: 54%  M: 55% |
|  | MKVILLEDVKGLGKKGDVVEVKDGYARNFLIPQGLAVEATKGNLKEL | | | |
| DHFR | 6195 | 3059 | T: 63 (1.0%)  M: 1884 (28.8%)  U: 1112 | T: 47%  M: 48% |
|  | MISLIVAVAENGVIGKDNDLPWHLPEDLKHFKALTMGHPVIMGRKTFESIGRPLPGRRNIVLTRDPDYQAEGAEVVHSLEEALALAKEAEEVFVIGGAEIYAQALPLADRLYLTEIDADFEGDTFFPEIDSEWKEVSREEHPADEKNGYDYTFVTYERKK | | | |
| AK | 2768 | 1445 | T: 159 (4.4%)  M: 835 (23.6%)  U: 451 | T: 56%  M: 57% |
|  | MRIILLGPPGAGKGTQAKRIVEKYGIPHISTGDMLRAAIKAGTELGKKAKSYMDAGELVPDEIVIGLVKERLAQPDCNGFLLDGFPRTIPQAEALDELLKELGVKLDAVIELDVPDEELVERISGRRVCPASCGRTYHVKFNPPKVEGVCDVCGEELIQRDDDNEETVRKRLEVYHEQTAPLIDYYKKKGLLVTVDGTGSIDEVFADILAALGKKK | | | |
| PGK | 4961 | 2361 | T: 234 (4.2%)  M: 1374 (24.6%)  U: 753 | T: 56%  M: 57% |
|  | MNKTIDDLDLKGKRVLVRVDFNVPLKDGKITDDTRIRAALPTIKYLLEKGAKVILMSHLGRPKGEVDPKFSLAPVAKRLSELLGKPVKFADDCVGEEAEAAVAALKPGEVLLLENLRFHKGEEKNDPEFAKKLASLGDVYVNDAFGTAHRAHASTVGVAKFLPAAAGFLMEKELEALGKALENPERPFVAILGGAKVSDKIGVIENLLDKVDKLIIGGGMANTFLKAQGYEVGKSLVEEDKLDTAKELLEKAKEKGVKIVLPVDVVVADEFSADAETKVVPVDEIPDDWMGLDIGPKTVELFAEAIKDAKTIVWNGPMGVFEFEPFAKGTKAVAKAIAEATGAFSIVGGGDTAAAVNKLGLADKFSHISTGGGASLEFLEGKELPGVAALEDK | | | |
| The numbers of bacterial and archeal sequences are as reported in Table S1. Sequence source organisms identified by BacDive indicates the number of sequences in each MSA whose source organisms could be in the BacDive database (10). T and M indicate the number of sequence parent organisms that were classified as thermophilic and mesophilic by the BacDive database. U indicates the number of sequence parent organisms that had no growth temperature classification or had ambiguous classifications. For each family, the sequence is the consensus obtained if thermophilic sequences are removed from the MSA. Residues that differ from the consensus sequence of the full MSA are highlighted in red. | | | | |
